# A common gene signature of the right ventricle in failing rat and human hearts

**DOI:** 10.1101/2023.05.16.540913

**Authors:** Liane Jurida, Sebastian Werner, Fabienne Knapp, Bernd Niemann, Ling Li, Dimitri Grün, Stefanie Wirth, Axel Weber, Knut Beuerlein, Christoph Liebetrau, Christoph B. Wiedenroth, Stefan Guth, Baktybek Kojonazarov, Leili Jafari, Norbert Weissmann, Stefan Günther, Thomas Braun, Susanne Rohrbach, Michael Kracht

**Author notes:** Leadx contact (Michael Kracht).

## Abstract

The molecular mechanisms of progressive right heart failure are incompletely understood. We systematically compared rat models of pulmonary artery or aortic banding to identify the transcriptomic changes that occur over months in the failing right versus left ventricle. Detailed bioinformatics analyses of 181 RNAseq datasets from cardiomyocytes or whole heart samples from these models, led to the identification of gene signatures, protein, and transcription factor networks specific to ventricles, compensated or decompensated disease states and type of heart failure. RNA-FISH approaches confirmed PAB-mediated regulation of key genes and revealed striking, spatially heterogeneous mRNA expression in the heart. Intersection of rat PAB-specific gene sets with 95 transcriptome data sets from human patients with chronic thromboembolic pulmonary hypertension led to the identification of more than 50 genes whose expression levels strongly correlated with the severity of right heart disease. Together, these data define a conserved, differentially regulated genetic network that coordinates progressive right heart failure in rats and humans.

**Highlights:**

- Side-by-side comparisons of RV or LV transcriptomes in the slowly failing rat heart
- Identification of RV-specific gene sets in heart hypertrophy versus heart failure
- Identification of RV gene sets correlating with severity of human CTEPH
- Development of a core gene signature characteristic for RV failure

## Introduction

Heart failure (HF) is a common clinical syndrome characterized by the progressive inability of the heart to pump sufficient blood volume to the body’s organs. HF is a major public health problem worldwide, with a prevalence of approximately 1-2% in developed and increasing incidence in developing countries (Braunwald, 2015, 2021; Ziaeian & Fonarow, 2016). In the United States, 6 million persons suffer from HF, with approximately 1 million new HF cases diagnosed annually (Bozkurt *et al*, 2021). Despite improvements in HF-related survival rates between 2000 and 2012, there has been a recent increase in mortality rates for all age and sex subgroups. As a result, the mortality rates after hospitalization for HF remain high, at approximately 20% to 25% at 1 year (Bozkurt *et al*., 2021).

Left heart failure (LHF), the more common form of HF, is characterized by the inability of the left ventricle (LV) to pump blood effectively to the body, leading to a buildup of fluid in the lungs and other tissues. This is typically due to conditions such as coronary artery disease / myocardial infarction, hypertension, or LV valve defects (Chahine & Alvey, 2022).

Right-sided heart failure (RHF) occurs when the right ventricle fails to pump blood effectively through the lungs, leading to congestion in the systemic circulation (Taverne *et al*, 2021). This can be caused by pulmonary hypertension (PH), chronic pulmonary disease, or left-sided heart failure (Arrigo *et al*, 2019; Sabbah *et al*, 2022). Despite some similarities between LHF and RHF, important differences in functions, morphology, (patho)physiology, molecular mechanisms, and clinical management of the two conditions have been described (Dini *et al*, 2022; Friedberg & Redington, 2014).

The right ventricle (RV) has a thinner myocardium, a lower contractile force, and a more compliant wall than the left ventricle (Haddad *et al*, 2008). These differences reflect the lower pressure load placed on the RV by the pulmonary circulation. Additionally, the RV has a higher proportion of fibrous tissue and fewer myocytes than the left ventricle, rendering it more susceptible to fibrosis and myocardial remodeling in response to stress (Voelkel *et al*, 2006).

While the RV has long been perceived as the “low pressure bystander” of the LV, which consists of largely the same cardiomyocytes as the LV, it is in fact derived from a different set of precursor cells during embryonic development and has a complex three-dimensional anatomy and a very distinct contraction pattern (Friedberg & Redington, 2014). These differences also offer possible explanations for the observation that treatments developed for LHF are often not effective in RHF (Borgdorff *et al*, 2013; van der Bom *et al*, 2013). Therefore, mechanisms of right ventricular failure and its detection, and more specific different functional and molecular responses to pressure versus volume overload are still incompletely understood (Taverne *et al*., 2021).

Advances in surgical, medical and device therapies have demonstrated the capacity of the heart to reverse, at least in part, the failing phenotype but more careful characterization of the molecular changes associated with this effect, in particular in the transcriptome and extracellular matrix, is necessary to define true myocardial recovery, in which a failing heart regains both normal function and molecular makeup (Kim *et al*, 2018).

Transcriptomic studies have contributed fundamentally to our knowledge on myocardial remodeling in animal models of acute LHF and on human LHF, but key HF genes reported have been often inconsistent between studies (Dobreva & Braun, 2015; Joshi *et al*, 2021; O’Meara *et al*, 2015; Raghow, 2016). To resolve some of these issues, a recent meta-analysis curated and uniformly processed 16 public transcriptomic studies of left ventricular samples from 263 healthy and 653 failing human hearts, collected during heart transplantation, left ventricular assist device implantation, or surgical ventricular restoration, to derive a consensus signature of LHF (Ramirez Flores *et al*, 2021).

Similar resources and datasets have not previously been available for RHF. In this study, we performed a systematic investigation of rat models of chronic RHF (pulmonary artery banding, PAB) or LHF (aortic banding, AOB) to uncover the transcriptomic changes that occur over months in the failing RV compared with the failing LV. Deep bioinformatics analysis of 181 RNAseq data sets derived from cardiomyocytes and whole heart samples of these models led to the identification of gene signatures and protein and transcription factor (TF) networks that are specific for ventricles, disease states, as well as the type of heart failure. Data obtained by orthogonal RNA fluorescence *in situ* hybridization (RNA-FISH) approaches confirmed PAB-mediated regulation of key genes and showed large spatial heterogeneity of mRNA expression in the heart. Intersecting the rat PAB-specific gene sets with 95 human transcriptomic data sets from patients with chronic thromboembolic pulmonary hypertension (CTEPH) before and after pulmonary endarterectomy (PEA) resulted in the identification of more than 50 genes whose expression levels strongly correlated with the severity of right heart disease in humans. Together, these data define a genetic network representing a core gene signature of the failing right ventricle that coordinates progressive RVF, and will serve as a rich resource for the identification of novel biomarkers and specific targets for the treatment of the diseased right heart.

## Results

### Comparative identification of transcriptomic changes in rat models of progressive left and right heart failure

Heart failure in human patients typically develops slowly over several months and there is limited understanding if this accompanied by chamber-specific molecular pathways in left compared with right ventricle failure (Taverne *et al*., 2021). To systematically study molecular changes in both conditions, we established rat models of chronic, progressive heart disease using aortic banding (AOB), to induce left ventricle hypertrophy and failure, or, pulmonary artery banding (PAB), to induce right ventricle hypertrophy and failure (Andersen *et al*, 2020).

Specifically, non-constricting clips were placed around the pulmonary artery (for PAB) or the aorta (for AOB) of 6 weeks old weanling rats (as exemplified in Supplementary Fig. S1A by µCT). As the animals grew, these clips became increasingly constrictive resulting in compensatory heart hypertrophy at week 14 (PAB, AOB) followed by decompensated heart failure at week 29 (PAB) or week 33 (AOB). As controls for disease-and age-specific changes in gene expression patterns, the experimental protocol included animals which were sham-operated and sacrificed at week 14 (sham 1, S1) or at week 33 (sham 2, S2) of the study as shown schematically in Fig. 1A.

**Fig.1.**
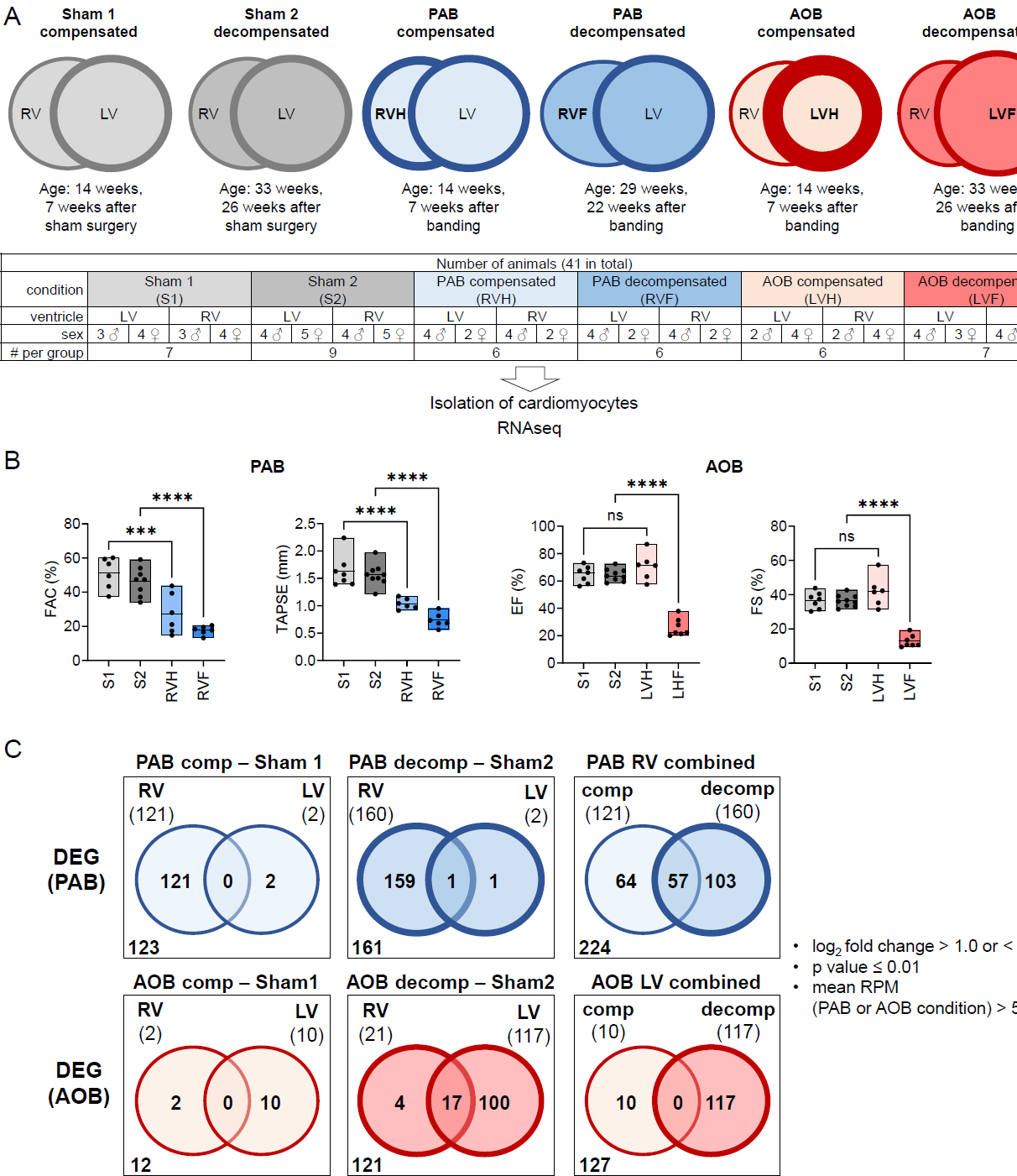
Identification of differentially regulated gene sets in rat models of progressive left and right heart failure. (A) Overview of animal study design. (B) Functional validation of compensated and decompensated states in response to pulmonary or aortic banding in the rat models of RHF and LHF. See also Fig. S1 for further functional data and the expression of PAB and AOB marker genes by RT-qPCR. FAC, fractional area change; TAPSE, tricuspid annular plane systolic excursion; EF, ejection fraction, FS, fractional shortening. (C) In total, 17341 rat genes were analysed by RNAseq. Based on twofold changes, mean read counts in disease conditions of > 50 and a p value ≤ 0.01, differentially expressed genes (DEGs) were identified in each condition by comparing the PAB or AOB groups to their corresponding sham groups. Venn diagrams indicate overlapping and distinct groups of DEGs for PAB (224 genes) and AOB (127 genes) that were further systematically analysed in this study. All data are provided in Source data for Fig. 1. See also Fig. S2 for the constitutively expressed rat transcriptomes in all conditions.

Disease progression was monitored clinically and by echocardiography. The assessment of RV (fractional area change, FAC; tricuspid annular plane systolic excursion, TAPSE) and LV parameters (ejection fraction, ES; fractional shortening, FS, and velocity time integral, VTI) confirmed that both, PAB and AOB resulted in terminal heart failure that proceeded through an intermediate compensatory state (Fig. 1B and Supplementary Fig. S1B). This was further validated by RT-qPCR analysis of whole heart samples which showed the expected increase in mRNA expression levels of three established marker genes for heart remodelling and failure (*Nppa*, *Nppb* and *Acta1*) in the affected ventricles (Supplementary Fig. S1C) (Stastna & Van Eyk, 2012; Stilli *et al*, 2006).

We isolated cardiomyocytes from the left and the right ventricles of all conditions from 6-9 animals per group (as shown in the table of Fig. 1A) and performed total RNAseq of all 82 samples.

Based on the mean read counts, altogether, 14170 (sham 1, LV), 14370 (sham 2, LV), 14220 (sham1, RV), and 14437 (sham 2, RV) expressed genes were detected in non-diseased conditions, of which 13627 (91 %) overlapped and defined the common gene sets of both, RV and LV cardiomyocytes (Supplementary Fig. S2A, B). Based on a twofold difference, a p value ≤ 0.01 and a minimal expression value of 50 reads per million mapped reads (RPM) per condition, we identified 230 differentially expressed genes (DEGs) between RV and LV in the sham 1 (S1) and 3120 DEGs between RV and LV in the sham 2 (S2) conditions, indicating that large part of the ventricle transcriptomes become differentially expressed in growing up animals (Supplementary Fig. S1C). These genes are involved in a plethora of biological processes that clearly differ between both ventricles, as shown by the clustered heatmap of the top 100 enriched pathway terms (Supplementary Fig. 1C). These results define the basal juvenile and adult chamber- and age-dependent transcriptomes of rat cardiomyocytes in healthy conditions (Supplementary Fig. S2A-D).

We then interrogated the RNAseq data sets to identify ventricle- and disease stage-specific changes in cardiomyocyte gene expression by comparing the PAB or AOB conditions to the corresponding sham controls (based on mean read counts in disease conditions > 50, twofold changes and a p value ≤ 0.01).

Ventricle-specific analysis of DEGs in the decompensated condition revealed 160 DEGs upon PAB in the RV and 117 DEGs upon AOB in the LV with only a few genes changing in the corresponding other ventricle (Fig. 1C, middle Venn diagrams). These data showed the banding model-specific transcriptomic response of the directly affected ventricle adjacent to the banded vessel.

Ventricle-specific analysis of genes in the compensated condition revealed a similarly large number of 121 genes that was specifically deregulated in the RV (but not the LV) upon PAB. In contrast, only 10 genes were deregulated in the LV upon AOB (Fig. 1C, left Venn diagrams). These data suggest that during compensatory hypertrophy, the RV reacts with a stronger gene-regulatory response.

This observation is also reflected in the overlap analysis of all DEGs of the RV or LV during PAB or AOB (Fig. 1C, right Venn diagrams). Upon PAB, twice as many genes (224 compared to 127) are deregulated in directly affected ventricles. In the PAB condition, only 25% (57 genes) are shared between the compensated and decompensated states (Fig. 1C, right Venn diagrams).

In conclusion, the direct comparison of two highly standardized rat models of chronic right or left heart failure clearly revealed disease- and ventricle-specific regulated gene sets and indicated that the transition from right heart hypertrophy to right heart failure is accompanied by a profound shift in gene expression programs.

### Different transcriptomic responses of left and right ventricles to PAB

Next, we focused on the 224 genes deregulated in cardiomyocytes under PAB conditions and examined their expression patterns in the RV compared with the LV during disease progression. Z-scaled expression values were used to cluster these genes into seven groups (clusters 1-7, Fig. 2A, B).

**Fig. 2.**
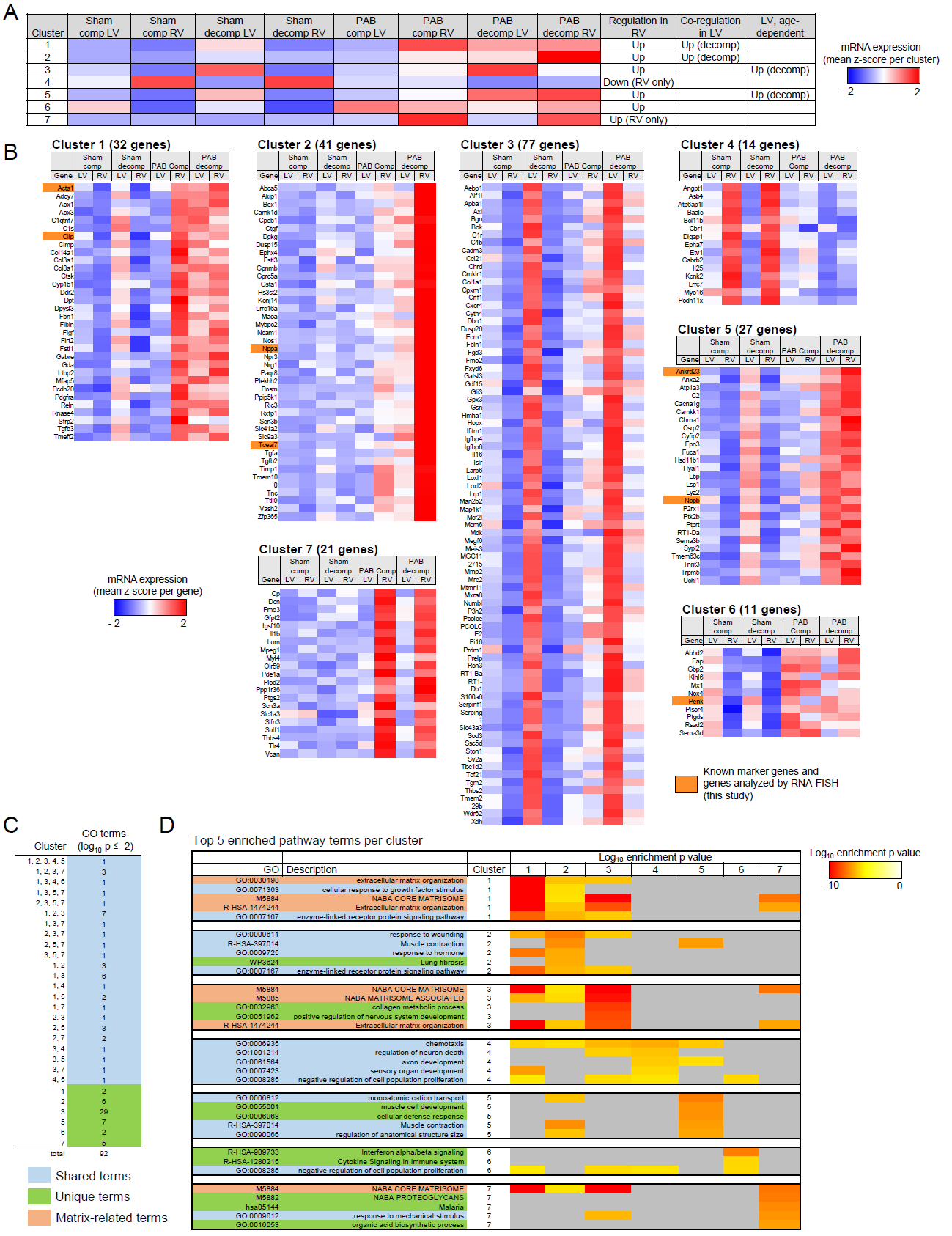
Different transcriptomic responses of left and right ventricles to PAB. (A) The mRNA expression values of all 224 PAB-regulated genes (see Fig. 1C) from right and left ventricles were hierarchically clustered by kmeans. The heatmap shows the averaged Z-score normalized read counts for all genes belonging to each of the seven clusters across all conditions in both left and right ventricles. The prevailing gene-regulatory phenotype is indicated in the three right columns. (B) Heatmaps of Z-scaled mean expression values for all individual 224 genes across the seven clusters and all conditions. Orange colors mark known (*Acta1*, *Cilp*, *Nppa*, *Nppb*) and new PAB-regulated genes that were highlighted in following analyses or further investigated throughout this study. (C) The genes of each cluster were examined for overrepresented pathway terms using Metascape (https://www.metascape.org) (Zhou *et al*., 2019) and multiple Venn analysis was performed to identify overlapping and distinct pathway terms for the gene sets of all 7 clusters. Only pathway term which enrichment p value ≤ −2 were considered. (D) The heatmap shows the top five enriched pathways for each cluster along with the corresponding enrichment p values for the other six clusters. Gray colors indicate that no enrichment was observed. All data are provided in Source data for Fig. 2.

As shown by the mean z-score of each cluster (Fig. 2A), but also at the level of individual genes (Fig. 2B), this analysis revealed seven distinct patterns of gene expression and regulation in the RV or LV according to age, RVH, and RVF.

Clusters 1-2 characterized PAB-induced genes that were strongly upregulated in the right ventricle in both the compensated and decompensated states (Fig. 2A, B). Only in decompensation were these gene sets also induced in the LV (Fig. 2A, B).

Cluster 3 gene sets were selectively upregulated in the LV of the oldest sham group already and thus comprise age-dependent LV-specific genes. On average, their expression was lower in the decompensating RV but still induced compared with the corresponding RV sham group (Fig. 2A, B).

Cluster 4 comprised the only set of genes that was more highly expressed in the RV (compared with the LV) in the healthy animals of the sham groups. These genes were downregulated under PAB conditions (Fig. 2A, B).

Genes from cluster 5 were strongly induced during decompensation in both the LV and RV. Similar to cluster 3 genes, their expression increased strongly only in the LV with age in the corresponding sham group (Fig. 2A, B).

Cluster 6 included genes that were strongly expressed only in the LV of the youngest sham group and whose expression decreased with age. These genes were already strongly induced in the compensated state in both the LV and RV under PAB conditions (Fig. 2A, B).

Finally, cluster 7 included the most specific PAB-induced genes, as their expression increased only in the RV during RVH and RVF (Fig. 2A, B).

The genes of the 7 clusters mapped to 92 pathway ontology terms, 51 of which were unique (Fig 2C, green font). For example, cluster 2 was enriched for genes involved in lung fibrosis (WP3624), while cluster 5 or 6 genes were annotated to muscle development (GO:0006968) or IFNα / β signalling (R-HSA-90973), respectively (Fig. 2D). Notably was the strong overrepresentation of pathway terms annotated to regulation of extracellular matrix in several gene clusters, such as clusters 1, 3 and 7 (Fig. 2D).

In summary, these data showed that many genes that are activated or repressed in the hypertrophied or failing RV are already highly expressed in the LV in an age-dependent manner and may become additionally co-regulated, especially in the decompensating heart. This also applies, for example, to already known marker genes such as *Acta1*, *Cilp*, *Nppa* or *Nppb* (highlighted in orange in Figure 2B). This implies that the particularly dynamic regulation of these genes in the RV may be masked by their expression in the LV when gene expression analysis is performed on the complete heart. At the functional level, the different expression patterns of these genes are most likely associated with a large variety of molecular remodeling processes and ventricle-specific functional changes that are coordinated at the transcriptome level under PAB conditions.

### Identification of gene-regulatory networks underlying the transition from right ventricular hypertrophy to failure

In the following, we restricted our analysis to the RV only to examine the time-dependent patterns of the 224 genes affected under PAB conditions during the transition from the healthy state (sham condition) to the compensatory hypertrophic (RVH) and decompensated (RVF) states. As before, Z-scaled expression values were used to cluster the 224 genes into five groups (clusters 1-5, Fig. 3A, B).

**Fig. 3.**
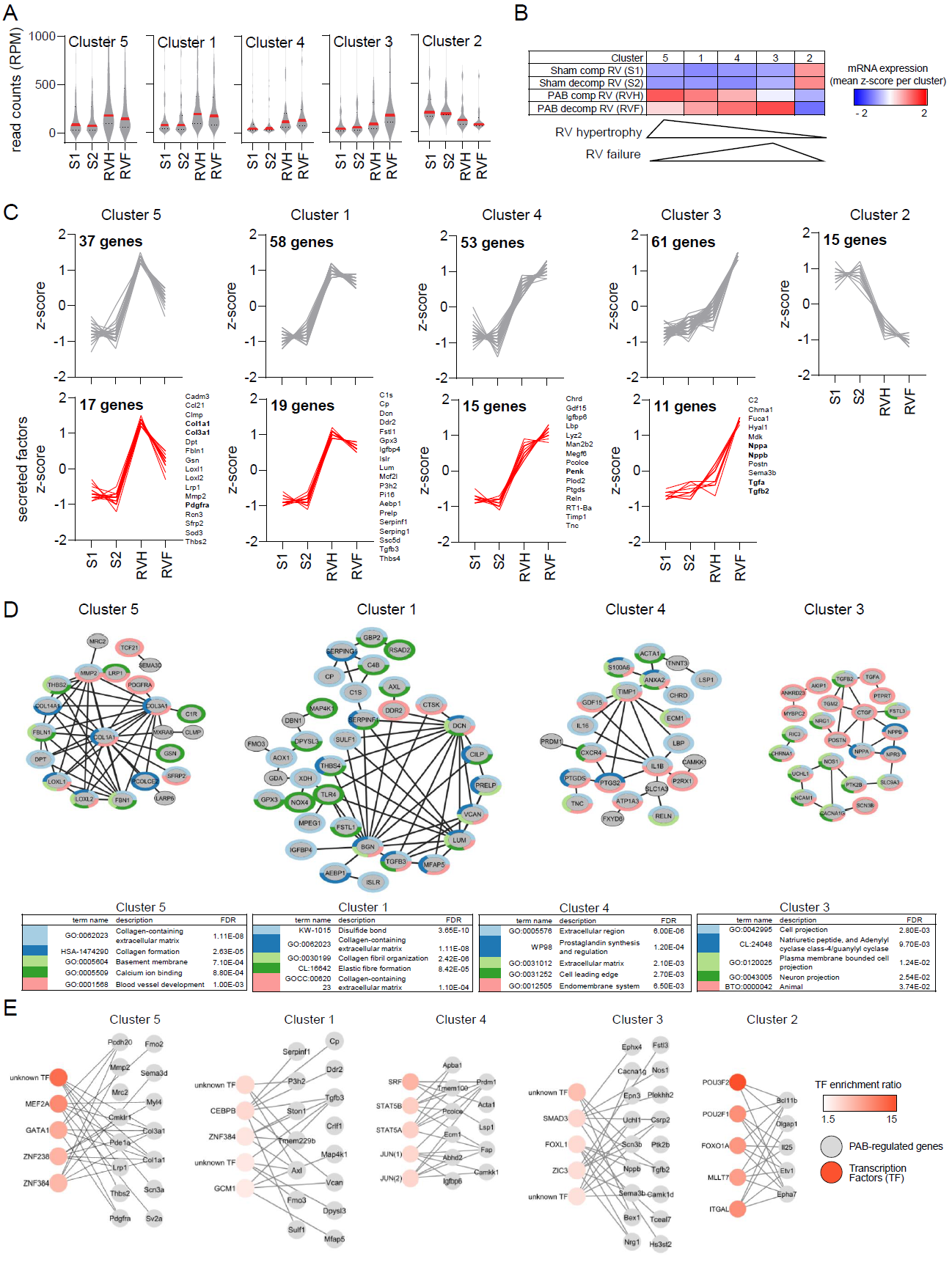
Identification of gene-regulatory networks underlying the transition from right ventricular hypertrophy to failure. (A) The mRNA expression values of all 224 PAB-regulated genes from right ventricles were segregated into five groups by hierarchical kmeans clustering. Violin plots show all normalized read counts (reads per million, RPM), medians (solid red lines) and 1^st^ and 3^rd^ quartiles (dotted red lines) (B) Z-scaled heatmap of mean mRNA expression for the genes of the five clusters from (A). Triangles indicate the overall regulation of cluster 5 and 3 which most specifically characterize the transition from the compensatory to the decompensated states. (C) Upper panels: Line graphs showing Z-scaled expression changes of all individual genes of each cluster. Lower panels: Red colours highlight the subgroups of genes from each cluster (also shown by gene name) that encode secreted factors according to a recent annotation of the secreted proteome by (Chen *et al*., 2019). The numbers of genes for each cluster are indicated in bold font. (D) Physical and functional protein networks for all genes (nodes) from each cluster based on STRING database annotations (Szklarczyk *et al*., 2019). All PPI categories of STRING (textmining, exeriments, databases, coexpression, neighbourhood, gene fusion and co-occurrence) and all pathway categories were selected. All shown edges had a STRING confidence core > 0.4. Shown are only genes (represented by gray nodes) which have at least one documented protein:protein interaction (PPI) as illustrated by gray edges connecting the nodes. Colors of node borders indicate the top five enriched pathway terms associated with each individual gene as determined by overrepresentation analysis using Cytoscape and the integrated STRING application. FDR, false discovery rate for pathway enrichment. (E) All genes of each cluster were examined for their overrepresentation in annotated transcription factor gene sets (TF) of the Molecular Signatures Database (MSigDB) (https://www.gsea-msigdb.org/gsea/msigdb) (Subramanian *et al*., 2005). Shown are the top 5 significantly enriched TFs (reddish nodes) which are connected to their target genes (grey nodes) by grey connecting edges. The complete lists of the top10 enriched unique TFs and all unique target genes are shown in Fig. S3A. The mRNA expression values for these TFs in the rat heart are shown in Fig. S3B. All data are provided in Source data for Fig. 3.

As illustrated by the normalized gene expression levels (Fig. 3A) and the mean Z-score of each cluster (Fig. 3B), these analyses revealed five distinct patterns of gene regulation specifically in the RV according to disease progression.

Cluster 5, 1 and 4 genes increased strongly upon PAB, whereby cluster 5 and 1 genes were highest at the compensatory state and decreased thereafter (Fig. 3A - C). In contrast, cluster 4 genes further increased upon decompensation (Fig. 3A - C).

Cluster 3 genes were not or not as strongly increased at the compensatory state, but were strongly induced in the failing RV (Fig. 3A - C).

Cluster 2 genes comprised the smallest group of genes which all were strongly downregulated during RVH and RVF (Fig. 3A -C)

Clusters 1,3,4,5 genes encoded 11 to 19 secreted proteins according to a recently published annotation of the secreted proteome (Chen *et al*, 2019). For example, cluster 4 contained *Penk* and *Timp1*, whereas cluster 3 included *Nppa*, *Nppb*, *TGFa*, and *TGFb3*. No transcript encoding a known secreted factor was observed in cluster 2 (Fig. 3C, lower graphs).

Many genes from clusters 1,3,4,5 encoded proteins that have known protein-protein interactions (PPI) and are parts of complex functional interaction networks as shown by the connecting edges of pathway nodes (Fig. 3D). Annotating individual network nodes with the five most strongly enriched pathway terms revealed a strong enrichment of multiple terms associated with collagen formation and extracellular matrix for clusters 1, 4 and 5 but also terms specific for one cluster such as prostaglandin metabolism (cluster 4) or calcium ion binding or blood vessel development (both associated with cluster 5) (Fig. 3D).

We then analysed if genes from clusters 1-5 were overrepresented in gene sets containing annotated bindings sites for specific transcription factors using the network transcription factor (TF) target functionality of WebGestalt (Wang *et al*, 2017). In total, this analysis identified 32 TFs regulating 106 of the 244 PAB genes (Fig. S3A). As shown at the examples of the top 5 most enriched TF, the genes of each cluster are predicted to be regulated by a specific combination of TFs, for example the cluster 5 genes *Col1a1*, *Col3a1*, *Pcdh20*, *Pdgfra*, *Scn3a*, *Sv2a* and *Cmklr1*are regulated by MEF2a, while the cluster 4 genes *Abhd2*, *Camkk1*, *Prdm1*, *Ecm1*, *Fap* and *Igfbp6* are regulated by JUN proteins (Fig. 3E). 17 of the predicted TF were found in the RV, of which *Jun*, *Srf*, *Nlf1*, *Mef2*, *Cebpb* and *Stat5b* were most strongly expressed, but overall their mRNA expression levels changed only weakly during PAB (Fig. S3B).

In summary, the transition phase from compensated to decompensated right heart failure is characterized by the activation of specific, relatively small gene regulatory networks encoding mainly protein networks involved in collagen and matrix metabolism. The time-dependent induction or repression of these genes correlates with different patterns of mRNAs encoding multiple secreted factors. These include known biomarkers such as *Nppa* or *Nppb*, which together with *Tgfa* or *Tgfb2* steadily increase in the failing RV (see cluster 3 genes), whereas cluster 5 genes such as *Col1a1*, *Co13A1*, or *Pdfgra* already decrease when the compensated situation shifts toward decompensation, suggesting that this group may serve as novel combination of biomarkers for early deterioration of RV function. Bioinformatic analyses also suggest that gene sets regulated in the same direction are under the control of specific TF networks whose activity preferentially occurs at the protein rather than mRNA level.

### PAB or AOB regulate largely different sets of genes in individual ventricles of the failing rat heart

So far, we focussed our analyses on mRNA expression changes in both ventricles in response to PAB. Using the same strategies, we addressed the question as to which extent the resulting sets of DEGs and their functions would overlap with genes regulated upon AOB.

The 127 genes that were specifically deregulated in compensated or decompensated AOB conditions (see Fig. 1C) were segregated into 4 clusters as shown by their Z-score normalized mean expression (Fig. 4A).

**Fig. 4.**
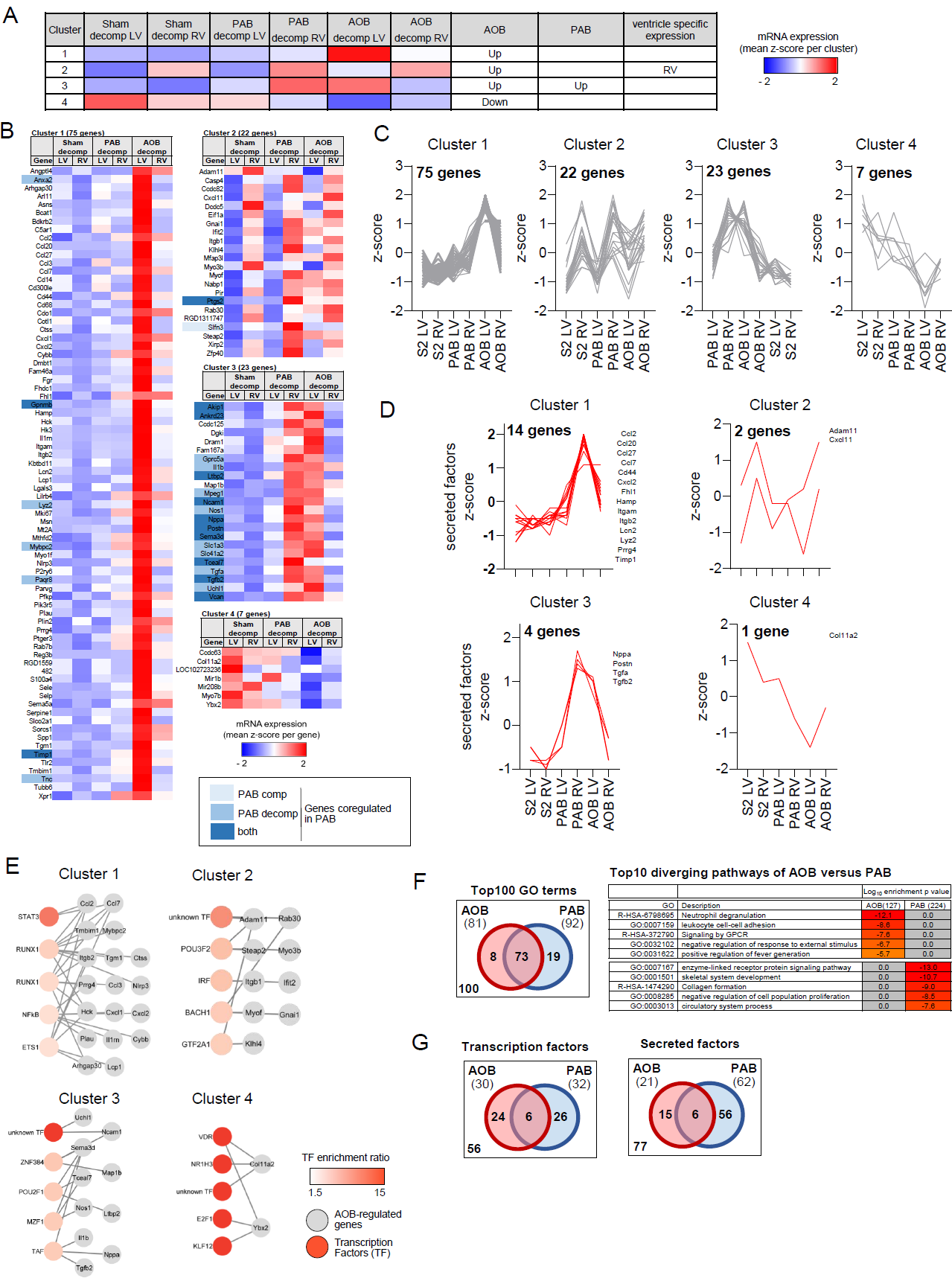
PAB or AOB conditions regulate largely different sets of genes in individual ventricles of the failing rat heart. (A) The mRNA expression values of all 127 AOB-regulated genes (see Fig. 1C) from right and left ventricles were segregated into four groups by hierarchical kmeans clustering. The heatmap shows the averaged Z-score normalized read counts for all genes belonging to each of the four clusters across all indicated conditions in both left and right ventricles. The prevailing gene-regulatory phenotype is indicated in the three right columns. (B) Heatmap of Z-scaled mean expression values for all individual 127 genes across the four clusters and all conditions. Blue colours mark PAB-regulated genes. (C) Line graphs showing Z-scaled expression changes of al individual genes of each cluster. The numbers of genes for each cluster are indicated in bold font. (D) Subgroups of genes from each cluster that encode secreted factors according to a recent annotation of the secreted proteome by (Chen *et al*., 2019). (E) All genes of each cluster were examined for their overrepresentation in annotated transcription factor gene sets (TF) of the MSigDB data base. Shown are the top 5 significantly enriched TFs (reddish nodes) which are connected to their target genes (grey nodes) by grey connecting edges. The complete lists of the top10 enriched unique TFs and all unique target genes are shown in Fig. S4A. (F) All DEGs from AOB (127 genes) or PAB (224 genes) were examined for overrepresented pathway terms. The Venn diagram show overlapping and distinct pathways for the top 100 most strongly enriched terms. The heatmap on the right shows the top 5 most strongly enriched terms that were unique for AOB or PAB. The complete lists of the top 100 pathway terms is provided a s clustered heatmap in Fig. S4B. (G) Venn diagrams demonstrating overlapping and distinct transcription and secreted factors identified in the DEG lists of PAB or AOB according to the analyses shown in Fig. 2-4. All data are provided in Source data for Fig. 4.

Cluster 1 contained 75 genes that were highly specifically induced in the LV only by AOB, of which only 7 genes (9%) were also regulated upon PAB (Fig. 4A-C).

Similarly, cluster 2 comprised a specific set of AOB-regulated genes that were on average weakly downregulated in the LV upon AOB, but were constitutively higher expressed in the RV compared to the LV (Fig. 4A-C).

In contrast, the regulation of cluster 3 genes largely overlapped between LV upon AOB and RV upon PAB, respectively. This set of genes contained *Nppa* / *Nppb*, *Il1b*, *Tceal7 Tgfa* and *Tgfb2*, that were amongst the marker genes that we followed throughout this study. Interestingly, the inducible expression of these genes was strictly restricted to the corresponding ventricle adjacent to the banded vessel, thus they increased in the LV upon AOB and in the RV upon PAB (Fig. 4A-C).

Lastly, cluster 4 genes were strongly downregulated by AOB in the LV and weakly downregulated in the RV upon AOB, but not regulated by PAB. These genes therefore represent an AOB-specific set that is negatively co-regulated in the LV and RV (Fig. 4A, B).

Altogether, only 27 (21%) of the 127 AOB genes were also co-regulated in both PAB conditions (Fig. 4A-C).

Cluster 1 genes contained 14 secreted factors, including four chemokines (*Ccl20*, *Ccl27*, *Ccl2* and *Cxcl2*) and two integrins (*Itgam*, *Itgb2*) which regulate leukocyte trafficking and activation (Hughes & Nibbs, 2018), while the other 3 clusters encoded only 7 secreted factors (Fig. 4D).

We identified 21 unique TFs suggested to regulate 64 AOB target genes of clusters 1-4, the top 5 of which are shown in Fig. 4E and supplementary Fig. S4A.

Pathway analysis showed that 27 of the Top100 pathway terms enriched for the 127 AOB or 224 PAB-regulated genes were different between AOB or PAB, while 73 terms overlapped (Fig. 4F and supplementary Fig. S4B). Three of the unique terms for AOB (R-HSA-6798695, GO:0007159, GO:0031622) referred to inflammatory processes, consistent with the regulation of secreted inflammatory mediators found in the highly LV AOB-specific cluster 1(Fig. 4F).

In conclusion, the direct comparison of the AOB with the PAB models indicated major differences between RHF and LHF that manifested at the gene, secreted factor, transcription factor network and pathway levels as shown by the Venn diagrams in Fig. 4F and Fig. 4G.

### Spatial regulation of PAB-regulated genes in the whole rat heart

To validate the DEGs regulated in the RV upon PAB, we followed two strategies. First, we reproduced the AOB and PAB animal studies, but this time subjected whole heart samples from the LV, RV and the septum to RNAseq, to test the possibility that the isolation procedure of cardiomyocytes contributed *per se* to the regulation of DEGs (Supplementary Fig. S5A). Second, we examined and quantified the *in situ* mRNA expression of prototypical PAB-regulated genes by single molecule (sm) RNA-FISH in whole heart sections (Fig. 5).

**Fig. 5.**
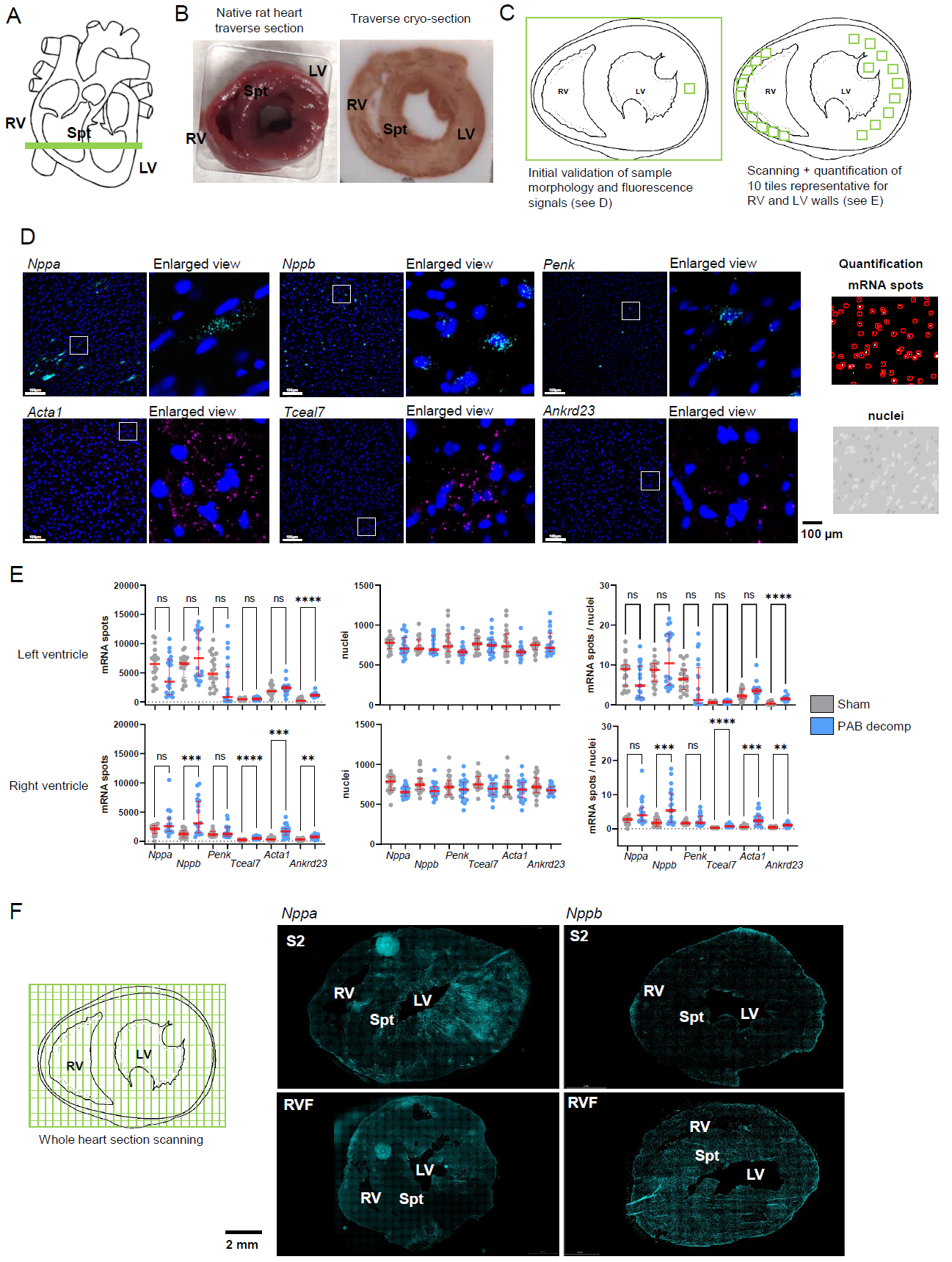
Spatial regulation of PAB-regulated genes in the whole rat heart revealed by RNA-FISH. (A) Scheme of the location of traverse sections of the heart used for smRNA-FISH. (B) Rat heart samples before and after processing for cryo-sections. (C) Schematic of the strategy for scanning the walls of the right and left ventricles through 10 tiles of equal-sized sections each. (D) Representative images of smRNA-FISH performed on 7 µm cryo-sections. Samples were hybridized with pairs of probes for the indicated transcripts and mRNAs were visualized by two different fluorophores. Nuclei were stained in parallel with DAPI. White inserts are shown as enlarged view on the right side of each image to demonstrate the spatial distribution of mRNA signals in the dense cardiomyocyte cell layers. Right upper and lower panels demonstrate the automated detection and quantification of mRNA spots (upper image) or nuclei (lower image) of each section by ICY software (https://icy.bioimageanalysis.org/) (de Chaumont *et al*, 2012). White scale bars indicate 100 µm. (E) Rat hearts from controls or PAB operated animals were obtained at the RHF states (sham 2, PAB decompensated) and processed for smRNA FISH as shown above. The graphs show quantification of mRNA spots, nuclei and mRNA spot signals normalized for cell number of each section according to the nuclei counts. Scatter plots show data from 20 sections derived from two biologically independent experiments. Red lines show medians, whiskers show 1^st^ and 3^rd^ quartiles. Significant changes were identified by one-way ANOVA; asterisks indicate p values (*p ≤ 0.05, **p ≤ 0.01, ***p ≤ 0.001, ****p ≤ 0.0001). (F) Left panel: schematic strategy for whole heart scans. Right panel: Exemplary images of the spatial distribution of Nppa and Nppb mRNA signals across the whole heart before and after PAB. The scale bar is 2000 µm. All data are provided in Source data for Fig. 5.

We filtered the whole heart data sets for significant changes (p ≤ 0.01) of DEGs in the RV of both PAB conditions. 47 out of 674 DEGs observed in whole heart samples overlapped with the principal set of 224 PAB-regulated genes of cardiomyocytes (Supplementary Fig. S5B). These genes mapped to a set of common pathways such as muscle contraction and development (R-HSA-397014, GO:0061061), circulatory system (GO:0003013) and blood vessel development (GO:0001568) (Supplementary Fig. S5C). These data also showed that overlapping genes were more strongly regulated in isolated cardiomyocytes compared to whole heart samples suggesting that whole heart sequencing, besides detecting genes from other cell types of the heart, reduces sensitivity for CM-specific DEGs as expected (Supplementary Fig. S5D).

From the 47 genes found by both approaches, we chose *Ankrd23*, *Tceal7*, *Penk*, *Nppb*, *Acta1* for follow-up smRNA-FISH because they were strongly regulated in the RV and showed high read counts in RNAseq analyses (improving the chance of detecting them by smRNA-FISH) as evidenced by the heat map shown in Supplementary Fig. S5D.

We prepared traverse cryo-sections of whole hearts from sham and decompensated PAB animals as shown schematically in Fig. 5A exemplarily in Fig. 5B. Samples were then simultaneously hybridized with pairs of probes for *Nppa*, *Nppb*, *Penk*, *Acta1*, *Ankrd23*, or *Tceal7* using two different fluorophores (C1, C2). Then, 10 representative sections of right and left ventricle walls were scanned (Fig. 5B-C). As shown by examples of original images, *Nppa*, *NppB*, *Penk* and *Ankrd23* mRNA spots showed a remarkably heterogeneous expression pattern across individual CMs, while *Acta1* and *Tceal7* mRNA spots were detected more evenly in many CMs (Fig. 1D). The cumulated numbers of individual mRNA spots and the counts of all nuclei from all 10 sections per heart region (RV, LV) were then collected from sham and decompensated PAB animals (Fig. 5E). For normalization, mRNA data were divided by the number of cell nuclei, which were detected by DAPI staining, and this parameter was used as a proxy for mRNA spots per cell (Fig. 5E). The quantification of mRNA spots from many thousands of CMs confirmed significant PAB-dependent increases in *NppB*, *Acta1*, *Tceal7* and *Ankrd23* only in the RV, whereas *Ankrd23* was the only gene significantly changing also in the LV (Fig. 5E). *Penk* mRNA also increased, but only in a part of all sections (Fig. 5E).

As evident from the scatter plots, the mRNA spot densities were heterogeneously distributed across individual tissue sections (Fig. 5 E), corroborating the observations on the single cell variability of gene expression in individual, even adjacent CMs, as shown by examples of images in Fig. 5D.

This impression was further confirmed by mounting whole images from 400 scans of heart sections to cover the whole heart area. In sham animals, *Nppa* appeared to be prevalently expressed in the LV, whereas upon RHF, its expression increased also in more areas of the RV. *NppB* was only sporadically found in the RV of sham animals, whereas its expression increased in both ventricles upon PAB. (Fig. 5E). Although lower resolution and non-quantitative, these scans re-inforce the notion of spatially restricted gene expression in the heart.

Together, whole heart RNAseq and RNA-FISH confirmed many of the PAB-regulated genes and further indicated that strong peaks of localized gene expression occur in both, healthy and in diseased hearts. The reason for this spatial control of CM gene expression is unclear, but obviously has major implications for the interpretation of gene expression results derived from bulk RNAseq (where heart areas of strong gene expression may become “diluted”) or from heart biopsies (where strong areas of gene expression may be missed).

### Rat PAB-regulated genes overlap with gene sets of human right heart disease

To analyse the conservation and relevance of the PAB-regulated gene sets for humans, we used 95 RNAseq data sets from patients suffering from chronic thromboembolic pulmonary hypertension (CTPEH). Right ventricular biopsies were obtained during thoracic surgery at base line (BL) and for a subset of 24 patients also from the septum (by right heart catheter) during follow up (FU), 12 months after pulmonary endarterectomy (Fig. 6A). According to European Society for Cardiology guide lines, disease severity of the patients was scored by multiple parameters (Humbert *et al*, 2023). Based on cardiac index, tricuspid annular plane systolic excursion (TAPSE) / systolic pulmonary arterial pressure (sPAP) und N-terminal pro-brain natriuretic peptide (NT-proBNP), the patients were stratified into low, intermediate and high one year mortality risk groups (Fig. 6B) and ranked for disease severity by equally weighting these parameters, so that higher ranks correlated with lower mortality ((Fig. 6C).

**Fig. 6.**
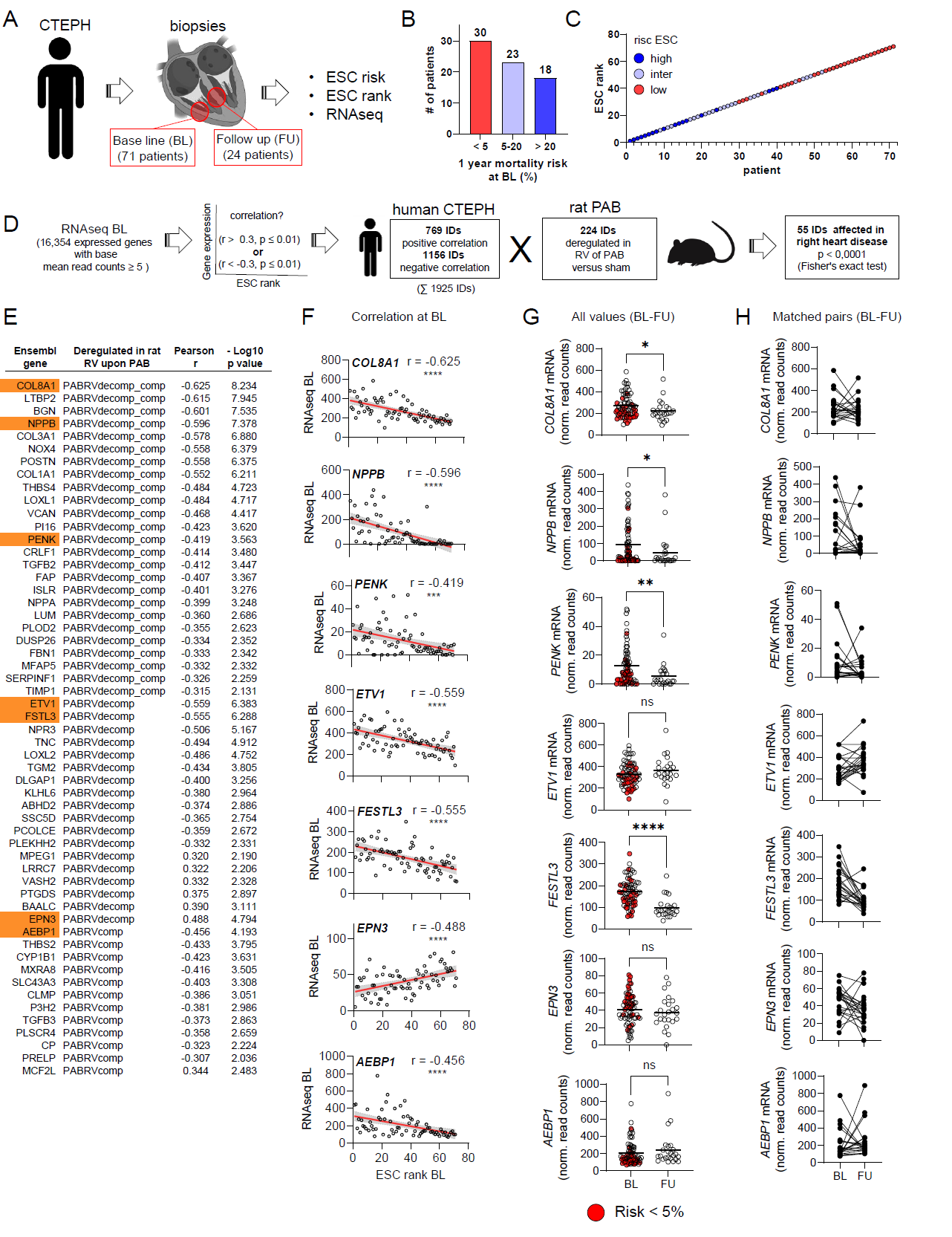
Rat PAB gene sets are deregulated in human chronic thromboembolic pulmonary hypertension (CTEPH patients) and their expression levels correlate with disease severity. (A) Schematic outline of patient cohorts and sample generation. At base line (BL), RNAseq was performed on 71 tissue samples taken from right ventricle walls during thoracic surgery of CTEPH patients. At follow-up (FU), RNAseq was also performed from 24 matching patient samples obtained by right heart catheter. In addition to RNAseq, clinical parameters were used to group patients at BL according to the disease risk (i.e. one year mortality, ESC risk) and to derive a disease severity rank (ESC rank) based on the European Society of Cardiology (ESC) criteria and guide lines for pulmonary hypertension (Humbert *et al*., 2023). (B) Graph showing the proportion of patients at BL with high, intermediate or low mortality risk. (C) Visualization of the relation of ESC rank to patient risk for all 71 patients at BL. (D) The expression values of all detected 16354 genes (IDs) with normalized read counts (reads per million, RPM) ≥ 5 from RV samples of each of the 71 human CTEPH patients at BL were correlated with ESC rank resulting in 1925 genes which showed a significant negative or positive correlation (based on Pearson r > or < 0.3 and a p value ≤ 0.01). This set was intersected with the 224 PAB-regulated genes from the rat model (see Fig. 1C), resulting in a significant (Fisher’s exact test, p < 0.0001) overlap of 55 genes. (E) Table showing all 55 genes overlapping between rat RVH (PABRVcomp) or rat RVF (PABRVdecomp) and human CTEPH at BL. The correlation coefficients and p values at BL which were determined as described in (D) are indicated. (F) Correlation of mRNA expression with ESC rank at BL for prototypical genes (marked in orange in D) Graphs show RPM for each of the individual patients on the Y-axis and ESC rank on the x-axis. Linear regression lines and 95% confidence intervals are shown in red and in gray colours, respectively. Graphs also include Pearson r and p values (*p ≤ 0.05, **p ≤ 0.01, ***p ≤ 0.001, ****p ≤ 0.0001). (G) Scatter dot plots showing mRNA expression (in RPM) of the genes form (F) at time of surgery (BL) and at follow up (FU). Red colors mark mRNA expression of BL samples from patients with lowest mortality risk. Black lines indicate means and asterisks indicate significant changes as obtained by Mann-Whitney test (*p ≤ 0.05, **p ≤ 0.01, ***p ≤ 0.001, ****p ≤ 0.0001). (H) Line plots of mRNA expression values for the genes shown in (F) and (G) at BL and FU for the 24 patients for which paired samples were available. All data are provided in Source data for Fig. 6.

Of the 16354 genes that were expressed at BL, 1925 correlated either negatively or positively with ESC rank, based on Pearson correlation coefficients > 0.3 and a p value of 1% (Fig. 6D). In this data set, 55 genes overlapped with 224-PAB-regulated genes (Fig. 6D-E, Source Data Fig. 6).

Most genes showed negative correlation, as exemplified by *COL8A1*, *NPPB*, *PENK*, *ETV1*, *FSTL3* and *AEBP1*, indicating that their levels increased with more severe RHF (Fig. 6E-F). Importantly, *COL8A1*, *LTBP2*, *NPPB* and *COLA3A1* were among the top 15 genes of all genes which negatively correlated with right heart disease of CTEPH patients at BL (Fig. 6E). Only 7 genes, such as *EPN3*, showed a positive correlation, indicating that their expression might be beneficial (Fig. 6E-F).

Quantification of mean changes and of pairwise comparisons between BL and FU confirmed a significant overall reduction of *COL8A1*, *NPPB*, *PENK* and *FSTL3* at FU (Fig. 6G). Accordingly, the mRNA levels of these genes obtained from patients at low risk were mostly below the mean of expression at BL (see red dots in Fig. 6G). This was not the case for *ETV1*, *EPN3*, *AEBP1* (Fig. 6G). Similarly, gene expression of 22 available paired samples showed a mixed pattern (Fig 6H). These results were in line with the interpretation that mRNA expression data obtained from two different regions of the heart (RV wall, septum) in a clinically heterogeneous group of patients are more variable and have limited sensitivity to reflect the course of disease.

A further comparison to human HF transcriptomic data was performed by intersecting the 224 rat PAB-regulated genes and all 2338 human CTEPH patients genes correlating significantly with ESC at BL (−log_10_ p values ≥ 2) with all significant genes extracted from 14 041 genes across 16 human LHF studies (−log_10_ meta analysis Benjamini-Hochberg corrected p values ≥ 1.3) (Ramirez Flores *et al*., 2021).

This analysis revealed that only 92 (2.2%) genes of all 4238 hLHF genes overlap with PAB-regulated genes, consistent with the notion that RHF and LHF are largely regulated by distinct transcriptomic responses, whereas the overlap was 679 (16%) between hCTEP and hLHF (Fig. 7A).

**Fig. 7.**
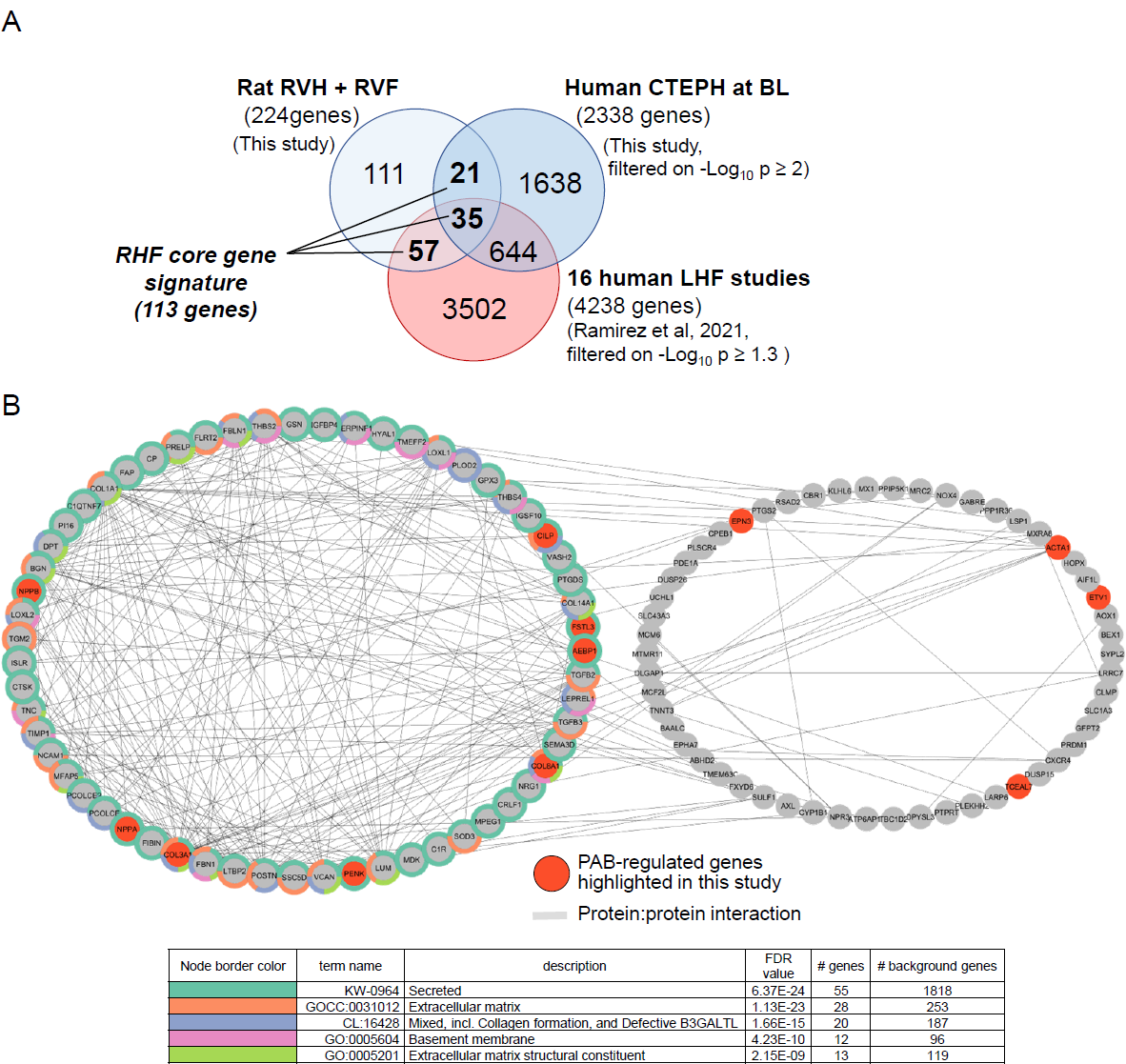
Meta-analysis of rat PAB, human CTEPH and human LHF gene sets to derive a core gene signature for right heart failure. (A) The 224 rat PAB-regulated genes were intersected with 2338 genes of human CTEPH patients correlating significantly with ESC at BL (p values on −log_10_ ≥ 2) and with all 4238 significant genes (based on −log_10_ meta analysis Benjamini-Hochberg p values ≥ 1.3) extracted from a total of 14041 genes across 16 human LHF studies (Ramirez Flores *et al*., 2021). The overlapping 113 genes constitute a core signature for RHF. (B) The 113 RHF core signature genes were examined for physical and functional protein networks for all genes (nodes) from each cluster based on STRING database annotations (Szklarczyk *et al*., 2019). All PPI categories of STRING (textmining, exeriments, databases, coexpression, neighbourhood, gene fusion and co-occurrence) and all pathway categories were selected. All shown edges had a STRING confidence core > 0.4. Colors of node borders indicate the top five enriched pathway terms associated with each individual gene as determined by overrepresentation analysis using the newest version of Cytoscape and the integrated STRING application (Shannon *et al*., 2003). FDR, false discovery rate for pathway enrichment. All data are provided in Source data for Fig. 7.

The total overlap of rat with human gene sets consisted of 113 genes, which contained all the genes that we consistently found to be regulated throughout this study by RVH or RVF in the rat (as marked with orange color in Fig. 7A). The fact that these genes were also found in the large hLHF data set is consistent with severe LHF also causing eventually RHF in some patients.

These 113 RHF-associated genes can be assigned to two large functional and physical protein interaction networks (Fig. 7B). The components of the large network show multiple interactions and are most highly enriched for secreted factors involved in the regulation of the extracelluar, collagen-containing matrix (Fig. 7B, left network). In contrast the components of the second network are not annotated to the top 5 enriched pathways and show fewer interactions, indicating that they have different functions and their roles in right heart disease still need to be defined (Fig. 7B, right network).

In conclusion, these data identified a unique set of RVH- and RVF-related genes that overlapped between the rat PAB models and human right heart disease. Many of these genes show a striking (mostly negative) correlation with disease severity in human patients. Therefore, these genes define, akin to the study by Ramirez-Florez et al. for LHF (Ramirez Flores *et al*., 2021), a first version of a RVF core gene signature.

## Discussion

Heart failure in humans usually manifests gradually over several months in response to (chronic) pressure or volume overload and other cardiac diseases. It is unclear whether the left and right ventricles share the same underlying pathophysiological mechanisms (Taverne *et al*., 2021). In this study, systematic comparisons of genes deregulated in rat PAB models with (i) AOB conditions, (ii) deregulated genes in CTEPH patients, and (iii) published meta-analyses of gene lists from LHF studies in patients led to the identification of a set of common genes and gene regulatory networks that we believe define a RVF core gene signature.

Our data show that cardiomyocytes from the RV and LV of the heart share more than 90 % of their transcripts. However, many of the RVF genes vary strongly and specifically in their expression levels in both, healthy conditions and in disease and in the following we discuss the potential roles of some prominent examples.

A predominant observation throughout our study is the altered mRNA expression of a large number of factors that alter the extracellular matrix (ECM) in the diseased RV. The myocardial ECM network contributes significantly to cardiac homeostasis by (i) providing structural support, (ii) facilitating force transmission and (iii) transducing molecular signals that regulate cell phenotype and function (reviewed in (Frangogiannis, 2019)).

In the failing RV, we found several genes encoding specific fibrillar type I and III collagens (*Col1A1*, *Col3A1*, *Col14A1*) amongst the most strongly regulated ECM factors in both, rat models and CTEPH patients.

Fibrillar collagens (types I, II, III, V, XI, XXIV and XXVII) function to provide three-dimensional frameworks for tissues and organs that confer mechanical strength as well as signalling and organizing functions through binding to cellular receptors and other components of the ECM (Bella & Hulmes, 2017).

Col I and Col III are abundant in the heart, whereby Col I is the predominant fibrillar collagen in adult mammalian hearts (Bashey *et al*, 1992; Hinderer & Schenke-Layland, 2019). The ratio of Col I to Col III in the left ventricles varies between mammalian species and with age (Medugorac, 1982). Several studies have shown changes in Col I and III levels in different models and types of left heart diseases (reviewed in (Frangogiannis, 2019)). Thus, both Col I and Col III have been reported to be overexpressed in pressure-overloaded hearts (Robert *et al*, 1994). In LV biopsies from human patients with aortic valve stenosis (AS) or regurgitation, comparable induction of Col I and III was noted and (in AS) positively correlated with LV end-diastolic pressure (LVEDP) and inversely with left ventricular ejection fraction (LVEF) (Fielitz *et al*, 2001). In a coronary artery MI model in rats, Wei *et al*. found a persistent increase of Col I levels (compared to a transient increase of Col III) in the non-infarcted areas of LV and RV and the RV post infarction, suggesting that Col I and III ratios are remodeled in response to hemodynamic stress (Wei *et al*, 1999). Based on the analysis of transmural biopsies from the LV free wall in patients with severe aortic stenosis with preserved ejection fraction and symptoms of heart failure, diastolic dysfunction was associated with increased nonmysial deposition of (predominantly) Col I collagen, resulting in increased extracellular matrix stiffness (Echegaray *et al*, 2017), corroborating results from an earlier study performed in high salt fed rats (Yamamoto *et al*, 2002). However, in that study, myocardial stiffening was associated with changes in collagen accumulation and enhanced collagen cross-linking rather than with the state of LV hypertrophy or LV failure (Yamamoto *et al*., 2002). Along these lines, in the hearts of spontaneously hypertensive rats (SHR), left ventricular end-diastolic (LVED) myocardial stiffness was not associated with changes in Col I:Col III ratio, but was likewise attributed to crosslinking (Norton *et al*, 1997). These observations suggest that the major fibrillary collagens play complex and diverging roles in different forms of cardiovascular and left heart disease.

Additionally, we found the non-fibrillar collagen gene *Col8A1* to be strongly regulated in both, rat models and CTEPH patients. Non-fibrillar collagens have been implicated in post-infarction remodeling (Shamhart & Meszaros, 2010). Double-deficient mice for the network-forming collagen type VIII (encoded by the *Col8A1* and *A2* genes) are viable, fertile, and show no major abnormalities (Hopfer *et al*, 2005). However, *Col8* KO mice exhibited increased mortality 3–9 days after aortic banding (AB) paralleled by increased LV dilatation from day one after AB. In the LV of *Col8* lacking mice, the expression of TGF-β1 was reduced. These data suggested that in acute LHF of mice, Col 8 promotes myofibroblast differentiation and fibrosis and protects from early mortality and LV dilatation in response to pressure overload (Skrbic *et al*, 2015). In a recent study, mice lacking *Col8* had reduced baseline blood pressure and altered ECM composition of carotid arteries, suggesting that Col 8 negatively regulates elastic fiber formation in the ECM matrix of blood vessels (Mohabeer *et al*, 2021).

We conclude that although the role of non-fibrillar components (basement membrane, proteoglycans, and glycoproteins) in cardiac fibrosis has been less extensively explored (Chute *et al*, 2019), there is emerging evidence that they are specifically deregulated in various forms of heart diseases, emphasizing the necessity to study their specific functional contribution also to RVF.

The cardiac ECM is not only a key regulator of cardiac remodeling through effects of its structural constituents on the mechanical properties of the heart, but also plays a major role in receptor-ligand regulated cellular responses (Frangogiannis, 2019).

In this study, we found the transforming growth factor (TGF)-β isoforms, *TGFb2* and *TGFb3*, amongst the core RV-regulated genes. TGF-β regulates a plethora of processes during tissue inflammation, repair, remodeling and fibrosis, including remodeling and fibrosis of the heart (Akhurst & Hata, 2012; Goletti & Gruson, 2015). The heart contains a significant amount of latent, pre-formed TGF-β in the ECM, which becomes released upon cardiac injury (Frangogiannis, 2014). The three highly homologous isoforms TGF-β1, TGF-β2 and TGF-β3 activate multiple cell types by binding to heteromeric complexes of transmembrane type I and type II serine/threonine kinase receptors (Akhurst & Hata, 2012). Evidence from mice lacking TGF-β1, 2 or TGF-β3 showed that although TGF-β isoforms share a receptor complex and activate the same principal canonical and non-canonical signalling machinery, they differ in their expression patterns and regulate specific, non-overlapping functions *in vivo* (Proetzel *et al*, 1995; Sanford *et al*, 1997). Isoform-specific functions of TGF-β in the heart are not well understood, because previous studies have largely focussed on TGF-β1 (Khan & Sheppard, 2006; Lim & Zhu, 2006; Liu *et al*, 2017; Xiao & Zhang, 2008). TGF-β was shown to activate expression of *Nox4*, another gene of the RV core signature, in cardiac fibroblasts and in models of airway inflammation (Rocic & Lucchesi, 2005; Zhang *et al*, 2019). Since Nox4 has been identified as a major source of oxidative stress in the failing heart, it will be interesting to study whether there is a functional link between Nox4, or recently discovered Nox4 isoforms relevant for human LHF, and *TGFb2* / *TGFb3* in RHF (Kuroda *et al*, 2010; Varga *et al*, 2017). Taken together, our data suggest that transcriptional *de novo* synthesis of TGF-β2 and 3 and activation of their downstream targets specifically contribute to RVF and these observations warrant further functional validation.

We conclude that it will be important to precisely dissect the mechanistic contributions of the highly regulated RV components of the ECM that have emerged from our bioinformatics analysis. In this context, it will be important to study their roles in (chronically) altered ECM turnover as a basis to better discriminate the pathogenetic basis of RHF from that of left heart diseases in both, animals models and human patients (Frangogiannis, 2019).

Another prevailing group of genes characteristic for RVF was annotated as secreted factors. The importance of “classical” secreted factors (such as prepro atrial natriuretic peptide (ANP), proANP, and fully processed ANP) in heart diseases is well recognized (Stastna & Van Eyk, 2012), as is the importance of paracrine factors released from cardiac progenitor cells (CPCs) to promote regenerative functions in heart diseases (Konemann *et al*, 2020; Samal *et al*, 2019).

Here, we consistently found upregulation of *Penk* mRNA in rat and human RHF. Penk (proenkephalin) encodes various endogenous opioid receptor agonistic peptides, called enkephalins, which have long been known to be highly expressed in the cells of the heart, including cardiomyocytes (Springhorn & Claycomb, 1992; van den Brink *et al*, 2003). The effects of opioids on the heart are complex and range from enhanced myocardial contraction to a (transient) decrease in heart rate and blood pressure (Bozkurt, 2019; van den Brink *et al*., 2003). In recent clinical studies of LHF patients with preseverd ejection fraction (LHFpEF), PENK plasma levels, used as stable surrogates for mature intracellular enkephalin synthesis, were prognostic for all-cause mortality and heart failure rehospitalisation (Kanagala et al, 2019). Higher PENK levels were also associated with more advanced LHF and glomerular and tubular dysfunction (Emmens et al, 2019). In the prevention of renal and vascular end-stage disease (PREVEND) study, in > 6000 participants, PENK levels were associated with both reduced renal function and higher N-terminal pro-B-type natriuretic peptide (NT-proBNP) levels, but were not independently associated with new-onset LHF, suggesting that PENK may serve primarily as a novel biomarker for renal glomerular dysfunction (Emmens *et al*, 2021). Based on these and other observations in LHF, a common PENK-dependent inter-organ pathway was postulated that affects both the heart and the kidneys (Matsue *et al*, 2017; Ng *et al*, 2017). In this pathway, high PENK levels might be detrimental, reflect a counter-regulatory response of an over-activated opioid system, or might be both, initially protective and later on maladaptive (Bozkurt *et al*., 2021; Emmens *et al*., 2021; Ng *et al*., 2017; Siong Chan *et al*, 2018). Almost nothing is known about a (specific) role of Penk and the opioid system in RVH and RVF, but the highly consistent upregulation of *Penk* mRNA in RHF observed in our study strongly suggests that its mechanistic role and potential use as an early biomarker for RHF should be analyzed in future studies.

### We also found multiple genes that had not been previously been implicated in RHF

*Tceal7*, a poorly characterized member of the transcription elongation factor A (SII)-like family of genes, was initially identified as a downregulated tumor suppressor in ovarian and other cancers (Chien *et al*, 2005). Mechanistic investigations revealed that TCEAL7 transcriptionally represses cyclin D1 expression and reduced levels of TCEAL7 promote DNA-binding activity of Myc and upregulation of Myc target genes (Chien *et al*, 2008). Additionally, *Tceal7* was found to be upregulated following cardiotoxin-mediated skeletal muscle injury in mice within 5 days, during the phase of muscle differentiation, and levels declined again thereafter in the following days (Shi & Garry, 2010). *Tceal7* was also identified as a direct, downstream target gene of myogenic regulatory factors (MRFs, e.g. Myod, Myf5, Myf6) and, collectively, available evidence suggested that Tceal7 repressed myogenic proliferation in favour of muscle cell differentiation (Sawada *et al*, 2022; Shi & Garry, 2010). A recent study further showed that transactivation of the *Tceal7* gene is mediated by a ternary complex of Mef2c, Creb1 and Myod assembling at the *Tceal7* promoter (Xiong *et al*, 2022).

The *Ankrd23* gene (encoding diabetes-related ankyrin repeat protein (DARP)) was isolated from hearts of diabetic mice and found to be highly expressed in heart, skeletal muscle and brown adipose tissues (Ikeda *et al*, 2003). Together with cardiac ankyrin repeat protein (CARP / Ankrd1) and ankyrin repeat domain protein 2 / ankyrin repeat protein with PEST and Proline-rich region (Ankrd2 / Arpp), Ankrd23 / DARP was classified as a member of stress-inducible muscle ankyrin repeat proteins (MARP) (Bang *et al*, 2014; Ling *et al*, 2017). The three MARP proteins are co-expressed in striated muscle tissues and share conserved motifs that interact with the giant filamentous titin polypeptide N2A segment to regulate stretch or stress responses (Miller *et al*, 2003). MARPs localize also to the nucleus and their expression is induced upon injury and hypertrophy (Ankrd1), stretch or denervation (Ankrd2), and during recovery following starvation (Ankrd23), suggesting that they are involved in (transcriptional) muscle stress response pathways (Miller *et al*., 2003). However, single, double and triple knockout mice of all MARPs are viable and no LV cardiac phenotypes were found in triple KO mice following 14 days of TAC (Bang *et al*., 2014).

The examples of *Penk*, *Tceal7* and *Ankrd23* highlight factors with some evidence for a function in muscle differentiation or LV heart diseases, which however, so far have not been investigated for their specific roles in right heart (patho)physiology.

The bioinformatics network analyses at the levels of TFs, pathways and protein interactions further suggest that the RVF core signature genes do not function in isolation but should rather be viewed as highly adapted gene-regulatory networks that are coordinated by specific patterns of secreted and transcriptional regulators and whose composition changes spatially and over time during the course of RVH and RVF. These data will, in combination with meta-analyses from studies assessing the (i) gene expression in individual cells types of the heart (Nicin *et al*, 2022; Tombor *et al*, 2021), (ii) spatially restricted expression patterns of genes (Calcagno *et al*, 2022; Longo *et al*, 2021), and (iii) protein levels in both ventricles (Chothani *et al*, 2019; Doll *et al*, 2017), greatly advance our knowledge on the molecular signatures of the RV in health and disease (Joshi *et al*., 2021).

In conclusion, this study identifies multiple, functionally connected genes that are consistently differentially regulated in chronic right heart failure of rats and humans. Though descriptive in nature, these data sets and their analyses provide a rich resource for future functional studies and the discovery of novel combinations of biomarkers that will serve to demarcate the transition from compensated to decompensated right heart failure.

### Limitations

The results and interpretation of this study are essentially based on extensive but descriptive RNAseq data sets. The postulated functional roles of individual genes or groups of genes require the performance of loss-of-function or gain-of-function experiments in the rat models under PAB or AOB conditions by depleting or overexpressing key factors in a ventricle-specific fashion, which were beyond our capabilities in the context of this study. The generation of a specified list of 113 RV core gene signature genes resulted from the adoption of relatively rigorously defined filtering criteria based on a combination of relative expression changes, expression levels, and T-test criteria in the rat PAB model. As with any bioinformatics analysis, it is clear that softening or tightening these filtering criteria would increase or decrease the number of disease-dependent regulated genes in the RV. Similarly, the annotation of DEGs as secreted, TF-regulated or protein network, is dependent on underlying databases and the parameters set. Here, we chose to use the default settings of Metascape, STRING, and WebGestalt, respectively. However, the publication of the entire raw dataset and all source data will allow colleagues in the field to (i) track all analysis results and (ii) generate alternative RV core signature lists on genes under their own chosen criteria.

## Methods

### AOB and PAB model in rats

Ascending aortic banding (AOB) and pulomary artery banding (PAB) were performed in 7 weeks old weanling hybrid rats (Wistar Kyoto x Lewis rats) by placement of a non-constricting clip around the vessels at the time of surgery (Weck Horizon Titanium Clip with an inner diameter of 0.6 mm for AOB and 1.1 mm for PAB, Teleflex Medical GmbH, Fellbach, Germany). Sham-operated animals served as age-matched controls. The surgery included an anterolateral thoracotomy under isoflurane anaesthesia in fully ventilated rats as described before (Knapp *et al*, 2020). As the animals grew up, the clip became constricting, resulting in a stage of compensatory LV (AOB model) or RV hypertrophy (PAB model) 7 weeks after surgery, followed by the onset of LV (AOB model) or RV (PAB model) failure 22-26 weeks after surgery. The clinical conditions of animals were characterized by two-dimensional and M-mode echocardiographic examinations in accordance with the criteria of the American Society of Echocardiography using a Vevo 2100 system (Fujifilm VisualSonics Inc., Toronto, Canada). Beside this evaluation of cardiac geometry and function, the animals were also inspected for clinical markers of heart failure, including tachypnoea, pleural effusion, enlarged liver and ascites. Both body weight and ventricular weight were recorded. All animal experiments conformed to the guidelines from Directive 2010/63/EU of the European Parliament on the protection of animals used for scientific purposes and were performed according to the regional authorities and ethics committees for animal research. The experiments were registered under the number G14-2017.

### In vivo Microscopic Computed Tomography (µCT)

Quantitative multiphase cine cardiac images were acquired using a Quantum GX µCT scanner (PerkinElmer, Inc, Waltham, MA) in conjunction with the contrast agent eXIA160XL (Binitio Biomedical, Inc, Ottawa, Ontario, Canada) as described previously (Kojonazarov *et al*, 2018).

Before imaging, the rat was restrained and an intravenous catheter prefilled with heparinized saline solution was introduced into a lateral tail vein. The rat was then placed into an induction chamber and anesthetized by inhalation of 3% isoflurane in oxygen. Afterwards the animal was placed on a scanner bed with a nose cone supplying isoflurane of 1.0% to 1.5% in oxygen. Two electrodes were placed on the right and left front paws of the rat to allow for continuous electrocardiographic monitoring. The contrast agent eXIA160XL (Binitio, Canada) was slowly infused at 5 µl / g of body weight at a rate of 0.3 ml / min with an infusion pump.

The scanner bed was translated longitudinally to align the animal chest within the center of the field of view. The scanner’s complementary metaloxide-semiconductor X-ray flat-panel detector was set to allow image acquisition with an X-ray tube voltage of 90 kV and current of 80 μA.

µCT data were collected in list-mode over a single complete gantry rotation with a total rotation time of 4 minutes (14 688 frames collected in total). Once the data were collected, the rat was removed from the scanner and allowed to recover fully from the anesthesia under supervision. Projection images acquired at 16-ms temporal resolution were transferred to a dedicated analysis workstation and reconstructed retrospectively. Reconstructed volumes were loaded into Analyze 12 software (Analyze Direct, Mayo Clinic). Short-axial image reformation was performed before LV and RV endocardial contour delineation, and the individual LV and RV volumes were calculated for all reconstructions. Fourier fitting was applied to reduce the fluctuation of the first derivative (Matlab, The MathWorks, Natick).

### Isolation of adult rat cardiomyocytes

Adult rat ventricular cardiomyocytes were isolated from rats as described in greater detail previously (Nippert *et al*, 2017). Briefly, hearts were excised under deep anesthesia, transferred rapidly to ice-cold saline, and mounted on the cannula of a Langendorff perfusion system. Hearts were perfused first in a non-re-circulating manner with 35 ml calcium-free perfusion buffer, then for 20-25 min in a re-circulating manner in a buffer supplemented with collagenase and 25 µmol / l CaCl_2_. Thereafter, the RV and the LV were separated. Afterwards both ventricles were minced separately and incubated for another 5 min. The remaining cell solution was filtered through a 200 µm nylon mesh. The suspension was centrifuged at 25 x g for 3 min to pellet down the RV and LV cardiomyocytes. The supernatant containing mostly the endothelial cells and fibroblasts was discarded and the cardiomyocytes were transferred into TRIzol™ reagent (ThermoFisher Scientific, Darmstadt, Germany) for RNA isolation according to the manufacturer’s instructions. Integrity and quality of the RNA was determined by automated electrophoresis using an Agilent 2100 Bioanalyzer system (Agilent Technologies).

### RNA isolation from whole heart tissue

Isolated ventricles were washed with ice-cold phosphate buffered saline and frozen in liquid nitrogen. The tissue was cut into small pieces and homogenized (rough surface glass douncer, 2ml, teflon pestle, Braun Melsungen) in 1ml TRIzol™ Reagent (ThermoFisher Scientific). After a second homogenization step in a fresh reaction tube using a plastic pestle, the samples were incubated for 5 min at room temperature. The insoluble material was removed by centrifugation (10 min, 12.000 x g, 4°C) and the supernatant was incubated again for 5 min at room temperature. 200 µl chloroform were added and after vortexing (15 sec) and further incubation for 3 min at room temperature, the samples were centrifuged (15 min, 12.000 x g, 4°C). The aqueous phase (containing the RNA) was transferred into a new reaction tube, mixed with 500 µl isopropanol and incubated for 10 min at room temperature. After centrifugation (10 min, 12.000 x g, 4°C), the supernatant was discarded and the RNA pellet was washed with 75% ethanol before air-drying and then dissolved in RNAse free water. Integrity and quality of the RNA was determined by automated electrophoresis using an Agilent 2100 Bioanalyzer system (Agilent Technologies).

### RT-qPCR

Trizol-purified total RNA was treated with rDNase using the Macherey-Nagel Nucleospin RNA Isolation kit according to the manufacturer’s protocol. Prior to cDNA synthesis, the RNA concentration was determined by measuring UV-absorption. mRNA measurements were performed by conventional RT-qPCR as described before (Jurida et al. 2015) using primer pairs for Nppa (5’-ttctccatcaccaagggctt-3’/5’-gcagatctatcggaggggtc-3’), Nppb (5’-ccggatccaggagagacttc-3’/5’-acaggatcacttgagaggtgg-3’) and Acta1 (5’-caatgagcgcttccgttgc −3’/5’-ctgcatgcggtcagcgata −3’). Each assay was performed in duplicate and validation of the correct PCR product was assessed by evaluation of the melting curve of the PCR products.

### Cardiomyocyte RNAseq

Ribosomal RNA depletion of total RNA isolated from cardiomyocytes, library preparation and deep sequencing was performed by Novogene on the Illumina Novaseq platform using 150 bp paired end set-ups. Raw data were processed using the Nextflow (Di Tommaso *et al*, 2017) and nf-core / RNAseq pipeline, version 3.2 (Ewels *et al*, 2020). Sequence reads were trimmed for adaptor / low-quality sequence using Trimgalore integrated into nf-core / RNAseq (parameter-quality limit: 20). Trimmed reads were mapped to Rnor6.0 using STAR integrated into nf-core / RNAaseq using the default parameters. Readcount extraction was perfomed using FeatureCounts (Rsubreads v2.2.6), disallowing multiple overlaps (Liao *et al*, 2014). The software STAR (Dobin *et al*, 2013) was used for alignment against the rat reference genome Rnor6.0 with UCSC feature annotations. Over 90% of all reads present in the 82 samples were mapped to genomic features of the rat genome and > 80% were uniquely mapped and represented by at least 25 million reads per sample. Differential expression was analyzed by DESeq2, version 1.28.1, using the independed filtering option and beta_prior LFC-shrinkage (Love *et al*, 2014), and the Galaxy platform (Galaxy, 2022) on the Galaxy environment of the Justus Liebig University Giessen.

### Whole heart tissue RNAseq

The polyadenylated RNA from whole heart tissue was enriched using the NEBNext® Poly(A) mRNA magnetic isolation module kit by adapting the manufacturer’s protocol to downscale the reaction volume to 33% (500 ng of total RNA in a volume of 34 µl with adapted amounts of beads and buffers). The RNAseq libraries were prepared using NEBNext® Ultra™ II directional RNA library prep kit for Illumina®, using a modified manufacturer’s protocol to reduce reaction volumes to 1 / 3 (7.7 µl mastermix added to RNA loaded beads). Libraries were quality controlled using Agilent Bioanalyzer high sensitivity DNA chips and DNA concentrations were determined using Qbit Analyzer with Qbit high sensitivity DNA reagent (Thermo Fisher scientific). Pooled libraries were prepared using SPRIselect beads (Beckman Coulter) before sequencing on an Illumina NextSeq 500 platform using 75 bp single end set-ups. Raw data were processed using the Nextflow (Di Tommaso *et al*., 2017) and nf-core / RNAseq pipeline, version 1.5 (Ewels *et al*., 2020). Sequence reads were trimmed for adaptor / low-quality sequence using Trimgalore integrated into nf-core / RNAseq (parameter - quality limit: 20). Trimmed reads were mapped to Rnor6.0 using STAR integrated into nf-core / RNAaseq using the default parameters. Readcount extraction was perfomed using FeatureCounts (Rsubreads v2.2.6) (Liao *et al*., 2014). The software STAR (Dobin *et al*., 2013) was used for alignment against the rat reference genome Rnor6.0 with UCSC feature annotations. For all 99 samples, over 95% of reads were mapped to the rat genome and > 75% were uniquely mapped. All samples reached a sequencing depth of 27 Million reads or above. Differential expression was analyzed by DESeq2, version 1.28.1, using the independed filtering option and beta_prior LFC-shrinkage (Love *et al*., 2014), and the Galaxy platform (Galaxy, 2022) on the Galaxy environment of the Justus Liebig University Giessen.

### RNA isolation and RNAseq analysis of samples from CTEPH patients

The present prospective study included a total number of 71 patients with a final diagnosis of chronic thromboembolic pulmonary hypertension (CTEPH), who were treated by pulmonary endarterectomy (PEA) at the Kerckhoff Heart and Thorax Center between 2016 and 2020. Biopsies of the free right ventricular wall from 71 patients were collected at base line (BL) during PEA. In a subgroup of 24 patients, RV myocardial biopsies were obtained during right heart catheterization (RHC) 12 months after PEA (follow-up, FU). In this case, to account for technical and safety aspects, the specimens were taken from the interventricular septum. Isolation of total RNA from both types of heart tissues was performed with Qiagen miRNeasy micro Kit and Covaris Cryo-Prep homogenization. 100 ng to 1 µg of total RNA was used for Hi-Mammalian whole transcriptome preparation (Takara Bio) and sequencing was performed on Nextseq2000 instrument with 1 x 72 bp single-end setup. Trimmomatic version 0.39 was employed to trim reads after a quality drop below a mean of Q15 in a window of 5 nucleotides and keeping only filtered reads longer than 15 nucleotides (Bolger *et al*, 2014). Reads were aligned versus Ensembl human genome version hg38 (Ensembl release 104) with STAR 2.7.10a (Dobin *et al*., 2013). Aligned reads were filtered to remove duplicates with Picard 2.27.1 (Picard Toolkit. 2019. Broad Institute, GitHub Repository. https://broadinstitute.github.io/picard/; Broad Institute; RRID:SCR_006525), multi-mapping events and ribosomal or mitochondrial reads. Gene counts were established with featureCounts 2.0.2 by aggregating reads overlapping exons on the correct strand excluding those overlapping multiple genes (Liao *et al*., 2014). The raw count matrix was normalized with DESeq2 version 1.30.1 (Love *et al*., 2014). Contrasts were created with DESeq2 based on the raw count matrix. Genes were classified as significantly differentially expressed at average count > 5, multiple testing adjusted p-value < 0.05, and −0.585 < log2FC > 0.585. The Ensemble annotation was enriched with UniProt data (Activities at the Universal Protein Resource (UniProt)).

### Classification of CTEPH cohort

Data on demographics, symptoms and comorbidities were collected for all 71 patients. Baseline characteristics concerning LV and RV dimensions, function and pressure gradients were further evaluated by transthoracic echocardiography and right heart catheterisation. All patients provided written informed consent for their participation in the study and approval of the institutional review board of the University of Giessen (AZ 44 / 14, 144 / 11, 145 / 11, 146 / 11, 199 / 15) was obtained. The investigation conforms to the principles outlined in the Declaration of Helsinki. Patients were grouped according to 1 year mortality risk by applying a modified version of the European Society of Cardiology (ESC) guidelines risk stratification model (Humbert *et al*., 2023), using the following criteria. Low risk (< 5%): cardiac index (CI) ≥ 2.0 L / min / m^2^, NT-proBNP < 300 ng / L, tricuspid annular plane systolic excursion (TAPSE) / systolic pulmonary arterial pressure (sPAP) > 0.32 mm / mmHg, intermediate risk (5-20%): NT-proBNP: 300-1100 ng / L, TAPSE / sPAP: 19 – 32 mm / mmHg and high risk (> 20%): CI < 2.0 L / min/m^2^, NT-proBNP > 1100 ng / L, TAPSE / sPAP < 0.19 mm/mmHg. Additionally, in the low or high groups 1 parameter was allowed to differ, but 2 out of 3 had to meet the pre-defined values. Otherwise, patients were assigned to the intermediate group.

### Single molecule RNA FISH

Isolated ventricles were washed with ice-cold phosphate buffered saline and frozen in liquid nitrogen. 7 µm tissue slices from adverse sections obtained from the two chamber ventricle level (without valves) were analysed by smRNA FISH using RNAscope fluorescent multiplex assay (Advanced Cell Diagnostics, Bio-techne.com) according to the manufacturer’s protocol for fresh frozen tissue. Briefly, tissue was permeabilized by dehydration and protease treatment, followed by hybridization of the selected RNA probes for *Nppa*, *Nppb*, *Penk*, *Acta1*, *Ankrd23* and *Tceal7* and amplification of the signals. Negative control probes for each channel (C1, C2) were directed against a bacterial RNA and provided by ACD Bio-techne.com. Microscopy analysis was performed using a Leica THUNDER imager (Leica Microsystems CMS GmbH) and Leica Application suite X (Version 3.7.4.23463), using pre-defined exposure times (PH - 55ms, DAPI - 80ms, C1 - 555nm - 100ms, C2 – 635nm – 700ms). After bright field preview scanning, the quality of morphology of the whole heart section and of fluorescence signals of single tiles were validated. Tile selection for region specific quantification were based on bright field scanning. Spots and nuclei (200 x magnification) were detected and quantified by Icy software (https://icy.bioimageanalysis.org) using the following settings: For nuclei quantification, the thresholder was adjusted to 15 k-means classes and HK-means to intensity classes of 12 and a minimum object size of 20 px. For spot detection, the object size was defined as 1 px with a sensitivity of 30. Spot number of each images were divided by the numbers of detected nuclei and normalized against background (factor: ratio of mean background level and background of the individual experiment). Imaging of all tiles across the entire section resulted in whole section overview images using Leica Application Suite X.

### Bioinformatic analysis and data visualization

All graphs and statistical tests (t-tests, ANOVA, correlation analyses) were performed using GraphPad Prism, version 9.4.1 (GraphPad Software, LLC) or Excel2016. Venn diagrams were designed with the help of Venn diagram tool (http://bioinformatics.psb.ugent.be/webtools/Venn/). For heatmap illustrations and cluster analysis Excel 2016 or the web tool Morpheus (https://software.broadinstitute.org/morpheus) were used, choosing a relative color scheme and k-means clustering with “one minus pearson correlation” based on row values. Protein-protein interaction network analysis were based on the newest version of the STRING database (https://string-db.org/) (Szklarczyk *et al*, 2019; Szklarczyk *et al*, 2023) and visualized using Cytoscape 3.9.1 using the integrated STRING functionalities and all ontology data bases of the STRING application (Shannon *et al*, 2003).

Pathway enrichment analyses were performed online by Metascape (www.metascape.org) using the following default settings: for each given gene list, pathway and process enrichment analysis were carried out with the following ontology sources: KEGG Pathway, GO Biological Processes, Reactome Gene Sets, Canonical Pathways, CORUM, WikiPathways, and PANTHER Pathway. All genes in the genome were used as the enrichment background. Terms with a p-value < 0.01, a minimum count of 3, and an enrichment factor > 1.5 (the enrichment factor is the ratio between the observed counts and the counts expected by chance) were collected and grouped into clusters based on their membership similarities. More specifically, p-values were calculated based on the cumulative hypergeometric distribution, and q-values were calculated using the Benjamini-Hochberg procedure to account for multiple testings. Kappa scores were used as the similarity metric when performing hierarchical clustering on the enriched terms, and sub-trees with a similarity of > 0.3 were considered a cluster. The most statistically significant term within a cluster were chosen to represent the cluster (Zhou *et al*, 2019). When multiple gene lists are provided, all lists are merged into one list called "_FINAL". A term may be found enriched in several individual gene lists and/or in the _FINAL gene list, and the best p-value among them is chosen as the final p-value. The pathway/process clusters that are found to be of interest (either shared or unique based on specific list enrichment) are used to prioritize the genes that fall into those clusters (membership is presented as 1/0 binary columns in the Excel spreadsheet). Note that individual gene lists containing more than 3000 genes are ignored during the enrichment analysis to avoid superficial terms; this is because long gene lists are often not random and generally trigger too many terms that are not of direct relevance to the biology under study (Zhou *et al*., 2019).

Enrichment analyses for transcription factor-regulated gene sets were performed online using WebGestalt (www.webgestalt.org) (Wang *et al*., 2017) and the transcription factor gene sets (TF) of the Molecular Signatures Database (MSigDB) (https://www.gsea-msigdb.org/gsea/msigdb) (Subramanian *et al*, 2005) with the follwing settings. Enrichment method: ORA, organism: rnorvegicus, enrichment categories: network_Transcription_Factor_target, ID type: genesymbol, reference list: all mapped entrezgene IDs from the selected platform genome. Parameters for the enrichment analysis: minimum number of IDs in the category: 5, maximum number of IDs in the category: 2000, FDR method: Benjamini-Hochberg, significance level: Top 10.

Secretome annotations were based on the published list of 6,943 high-confidence human secreted proteins which were generated from 330,427 human proteins derived from databases of UniProt, Ensembl, AceView, and RefSeq SPRomeDB (www.unimd.org/SPRomeDB). 6,267 of 6,943 (90.3%) of these proteins have the supporting evidences from a large amount of mass spectrometry (MS) and RNA-seq data as published in (Chen *et al*., 2019).

## Data availability

The rat RNAseq data sets of this study have been submitted to GEO: GSE216263 (https://www.ncbi.nlm.nih.gov/geo/query/acc.cgi?acc=GSE216263) and GSE216264 (https://www.ncbi.nlm.nih.gov/geo/query/acc.cgi?acc=GSE216264).

The remaining data generated in this study are provided in the Supplementary Information / Source Data sections. Source data are provided with this paper.

## Author contributions

L.J. designed, performed and analysed RNA-FISH experiments, Se.W. performed and analysed rat RNAseq experiments, B.N. performed all surgical interventions and echocardiography in rats, F.K., L.L. and S.R. performed all PAB and AOB animal experiments and isolated CM and RNA from CM for RNAseq, D.G. analysed human CTEPH patient data, St.W. performed RNA-FISH, A.W. analysed RNAseq data, K.B. analysed RNA-FISH data, C.L., C.B.W., S.G., L.J. and N.W. conceived and performed studies and sample acquisition from hCTEPH patients, S.G. and T.B. performed RNA seq analysis of hCTEPH patient samples, L.J. and M.K. performed all bioinformatics analyses and prepared all figures, M.K. conceived the study and wrote the initial complete draft, all authors contributed to the final version.

## Acknowledgments

This work was supported by the following grants from the Deutsche Forschungsgemeinschaft (DFG, German Research Foundation): SFB1213/2 (B03 [to M.K., S.R.], CP01 [to C.L.], CP02 [to B.K.], project 268555672); KR1143/9-2 (KFO309, P3, project 284237345); SFB1021/3 (C02 [to M.K.], Z03 [to M.K., U.L], project 197785619); and GRK2573 (RP5 [to M.K.], project 416910386). Work in the laboratories of M.K., N.W. and T.B. is also supported by the IMPRS program of the Max Planck Society and the Excellence Cluster CardioPulmonary Institute (EXC 2026: Cardio-Pulmonary Institute (CPI), project 390649896) and the DZL/UGMLC/ILH program.

We thank Oliver Dittrich-Breiholz, Research Core Unit Genomics, Institute of Physiological Chemistry, Medical School Hannover, Hannover, Germany for RNA sequencing of whole heart samples.

We thank Eckart Mayer, Kerckhoff Heart and Lung Center, Department of Thoracic Surgery, 61231 Bad Nauheim, Germany, for contributions to obtain biopsies from CTEPH patients.

## Supplementary figures

**Fig. S1.**
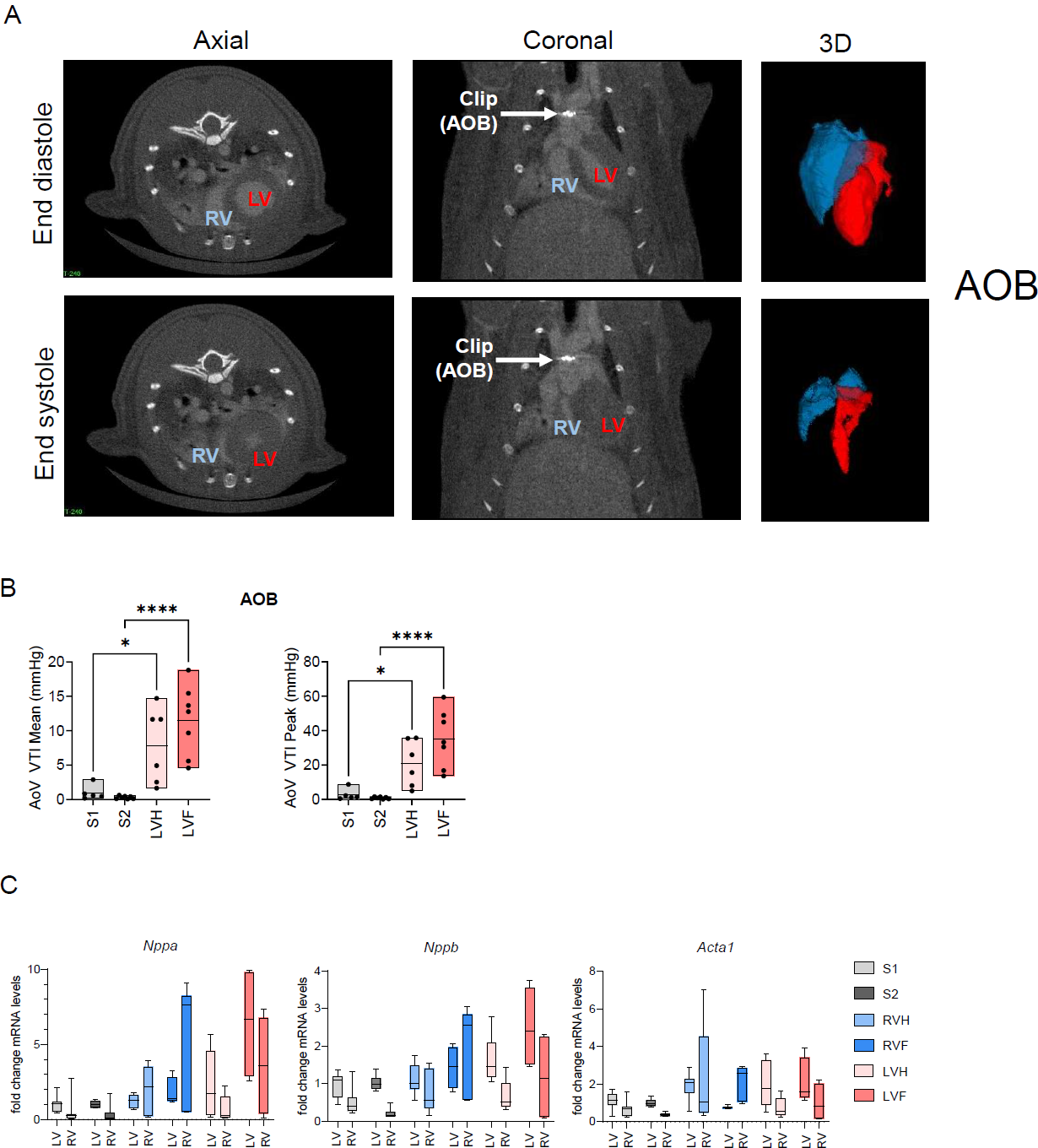
related to Fig. 1 Validation of HF marker genes in AOB or PAB conditions. (A) µCT images with contrast agent visualizing the position of the clip upon AOB of rats in the compensated, LVH state. (B) Functional validation of compensated and decompensated states in response to aortic banding in the rat model of LHF. VTI, velocity time integral. (C) Total RNA isolated from individual ventricles of animals treated according to the scheme shown in Fig. 1A was used to determine the expression of Nppa, Nppb and Acta1 by RT-qPCR. Relative mRNA levels compared to the mean of the Sham 1 condition of the LV are shown. Boxes of graphs show 1^st^ to 3^rd^ quartiles and medians. Data represent 14 (S1), 10 (S2), 14 (PAB comp), 6 (PAB decomp), 14 (AOB comp) and 8 (AOB decomp) animals. All data are provided in Source data for Fig. S1.

**Fig. S2.**
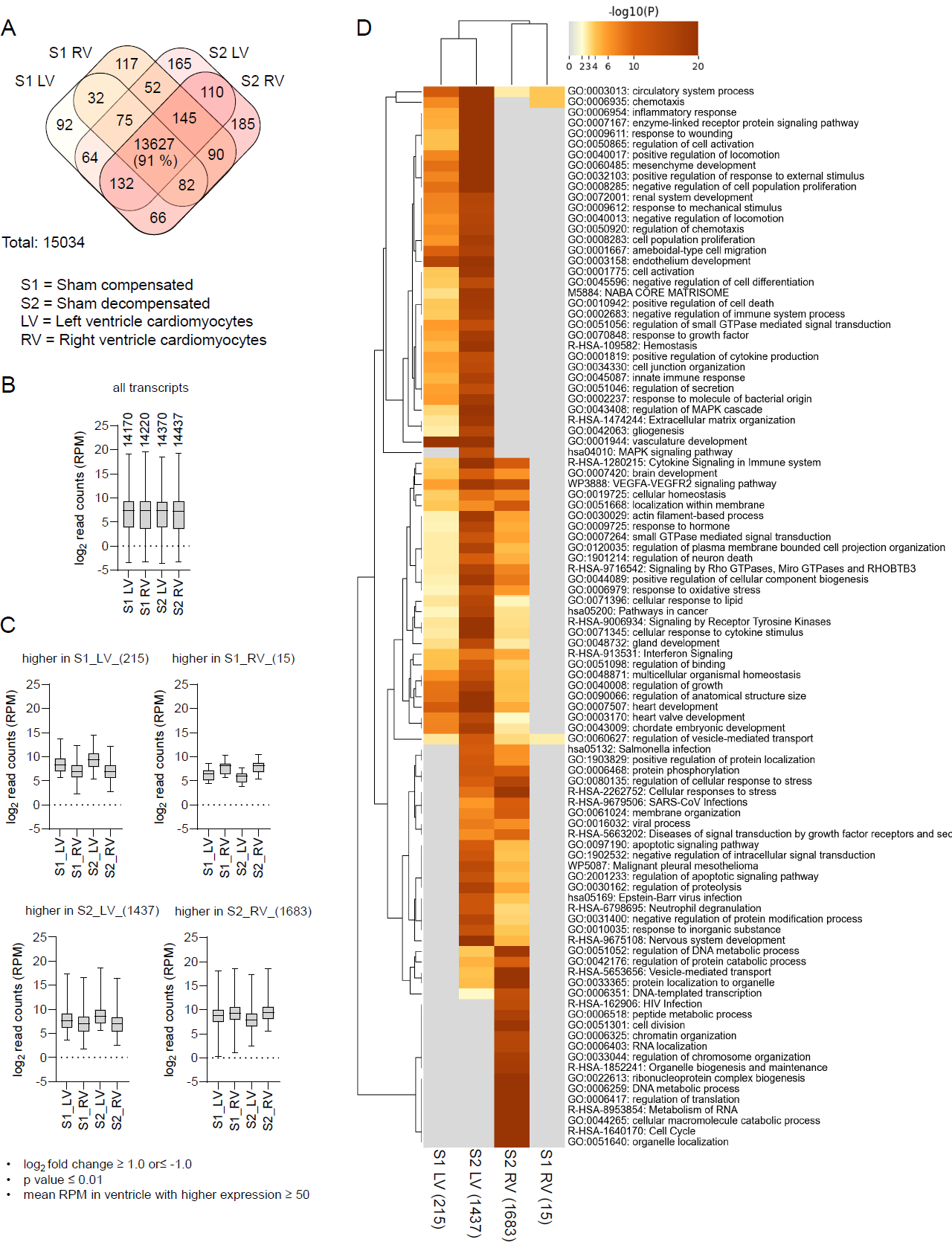
related to Fig. 1 Common and ventricle-specific sets of genes in healthy rat hearts at sham 1 and sham 2 conditions. (A) Venn diagram showing the number of overlapping and distinct sets of genes in left (LV) or right ventricle (RV) of young (Sham 1, S1) and older (Sham 2, S2) rats based on normalized mean read counts (reads per million, RPM). (B) Graph showing the distribution of mRNA expression levels of all assessed genes (Boxes show 1^st^ to 3^rd^ quartile, whiskers show minimum to maximum of all values). (C) Summary of all genes with differential expression between LV or RV based on a log_2_ fold change of ≥ 1.0 or ≤ −1, a p value ≤ 0.01 and mean RPM ≥ 50 in the ventricle with higher expression. Boxes show 1^st^ to 3^rd^ quartile, whiskers show minimum to maximum of all values. (D) Clustered heat map showing the top 100 overrepresented pathway terms for all genes with differential expression between LV or RV. All data are provided in Source data for Fig. S2.

**Fig. S3.**
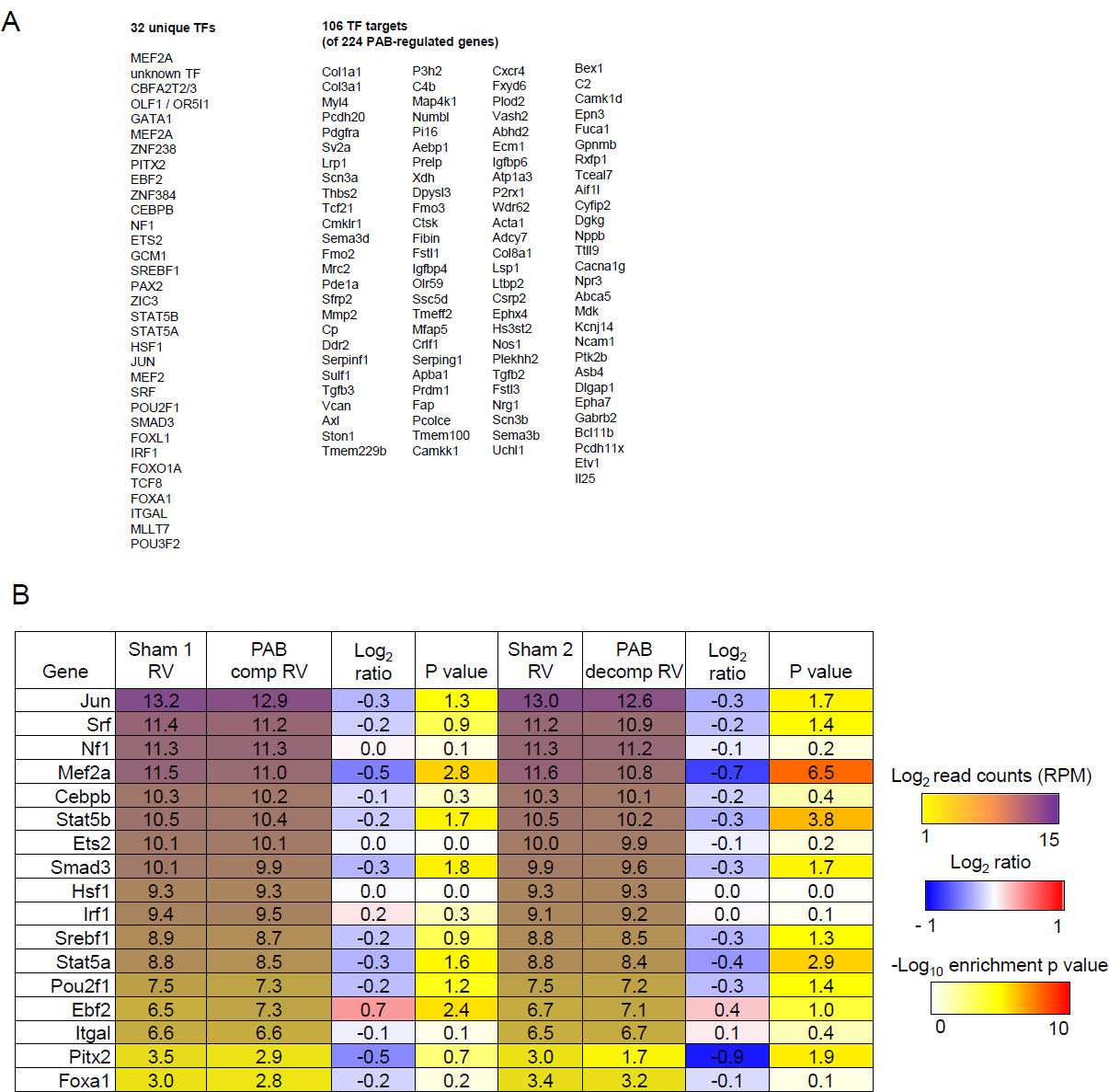
related to Fig. 3 Transcription factors (TFs) regulating PAB target genes. (A) Top 10 unique TFs and their targets genes. (B) Out 32 TFs, 17 TFs are expressed in the RV of S1 or PB animals. The heatmap shows read counts, fold changes and their significance, All data are provided in Source data for Fig. S3.

**Fig. S4.**
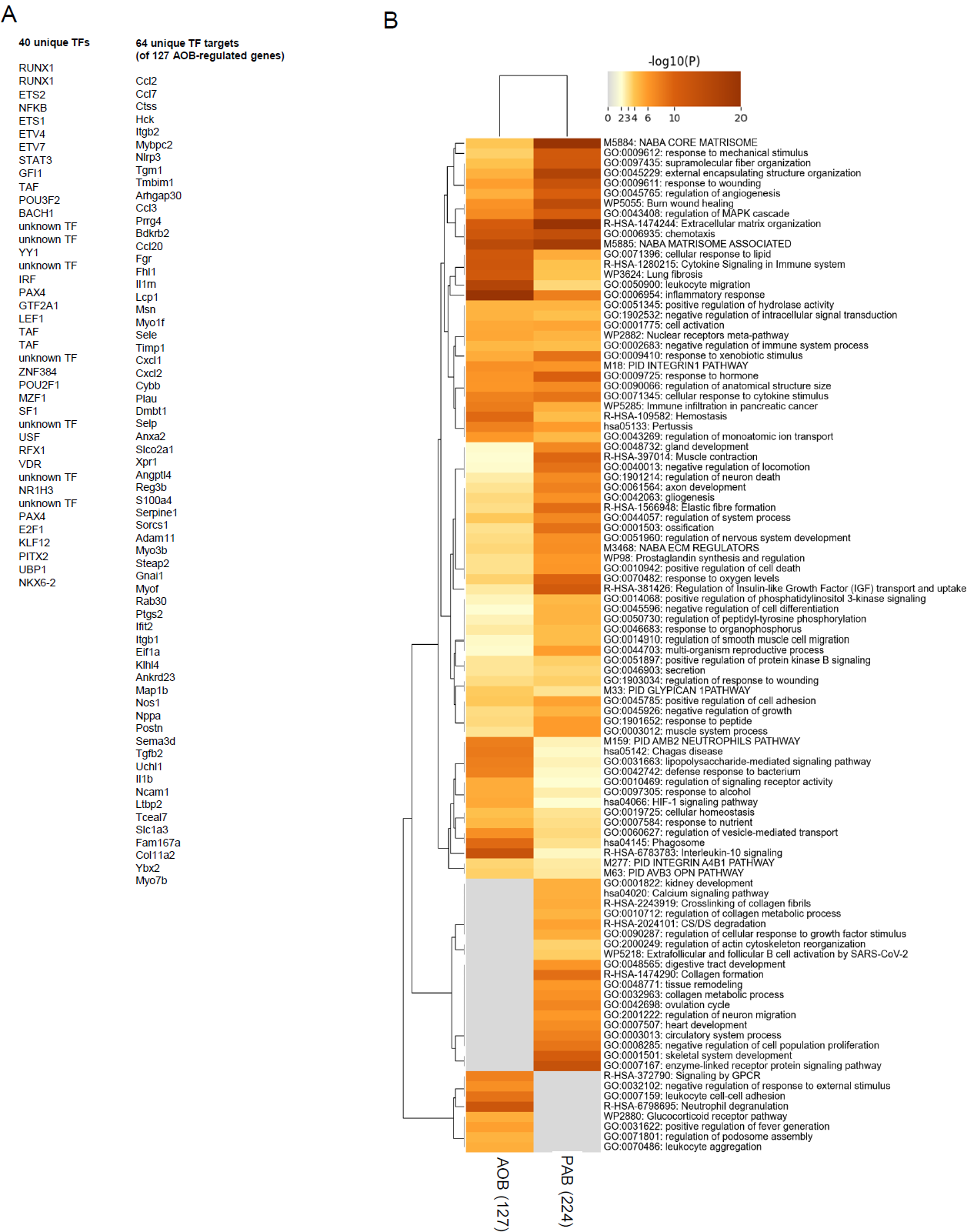
related to Fig. 4 Transcription factors (TFs) regulating AOB target genes and top 100 pathways terms associated with AOB or PAB. (A) Top 10 unique TFs and their targets genes. (B) The gene ontologies of all 224 PAB and 127 AOB target genes were determined by pathway overrepresentation analysis. The clustered heatmap shows the top 100 enriched pathway terms. All data are provided in Source data for Fig. S4.

**Fig. S5.**
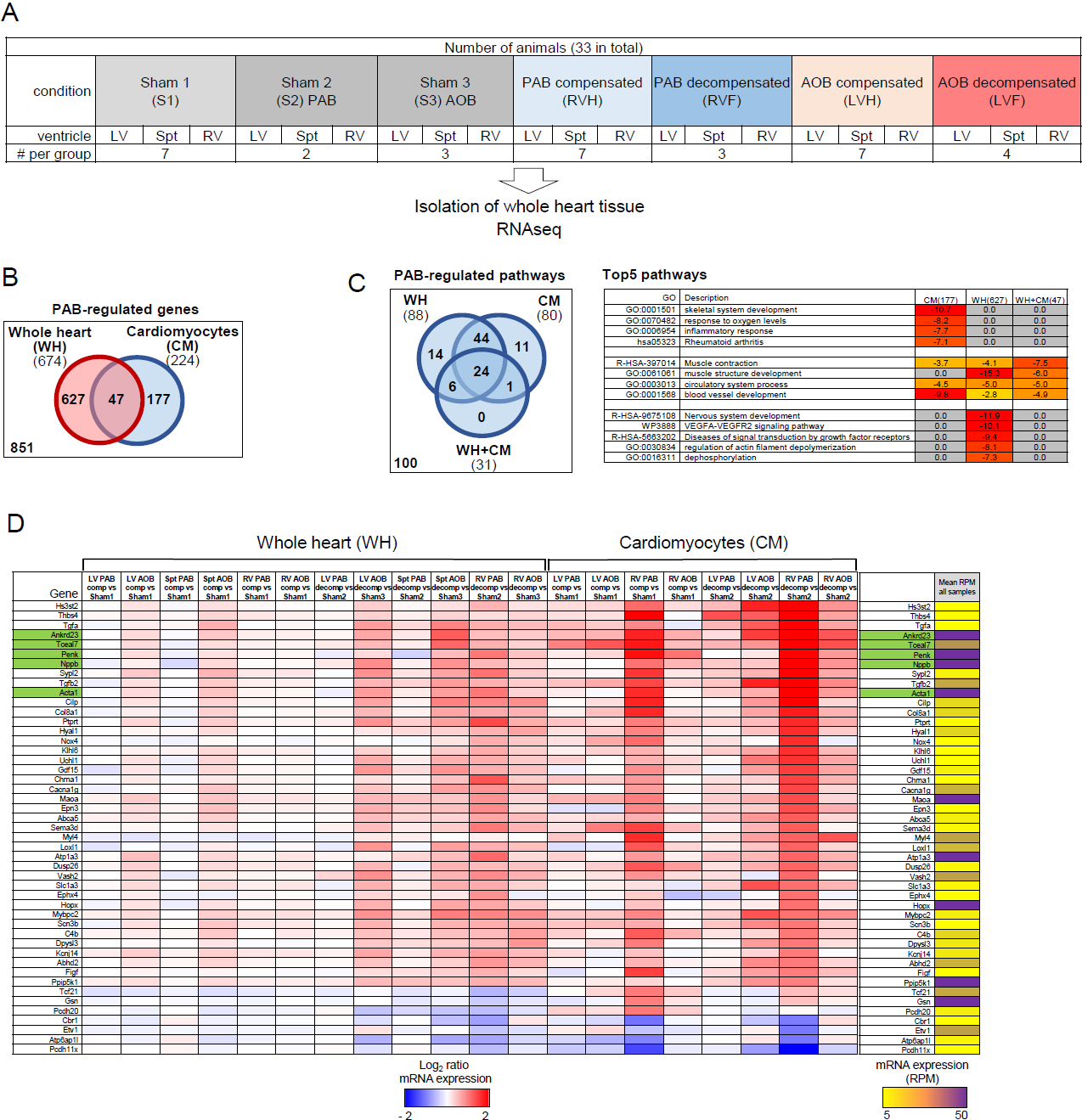
related to Fig. 5 PAB or AOB regulated genes in whole hearts compared to isolated cardiomyocytes. (A) Overview of animal study design. (B) Whole heart (WH) and cardiomyocyte (CM) RNAseq data sets were normalized together, joined and differentially expressed genes were obtained by Deseq2 analysis. Venn diagram showing the overlap of the 224 PAB regulated genes (based on the filter criteria shown in Fig. 1C) with 674 PAB-regulated genes obtained from whole heart RNAseq based on significant changes (p ≤ 0.01) of DEGs in the RV. (C) All DEGs from whole heart (674 genes) or cardiomyocytes (224 genes) as described in (B) were examined for overrepresented pathway terms using Metascape (Zhou *et al*., 2019). The Venn diagram show overlapping and distinct pathways for the top 100 most strongly enriched terms. The heatmap on the right shows the top 5 most strongly enriched common or unique terms. The complete lists of the top 100 pathway terms is provided in the source data for Fig. S5. (D) Heatmap showing fold changes across all conditions as described in (A) and mean overall expression values for all 47 genes jointly regulated in the whole heart or cardiomyocyte samples as described in (B). All data are provided in Source data for Fig. S5.

